# EPOP Restricts PRC2.1 Targeting to Chromatin by Directly Modulating Enzyme Complex Dimerization

**DOI:** 10.1101/2024.09.10.612337

**Authors:** Lihu Gong, Xiuli Liu, Xin Yang, Ze Yu, Siming Chen, Chao Xing, Xin Liu

**Author notes:** These authors contributed equally (Lihu Gong, Xiuli Liu, Xin Yang, Ze Yu).

## Abstract

Polycomb repressive complex 2 (PRC2) mediates developmental gene repression as two classes of holocomplexes, PRC2.1 and PRC2.2. EPOP is an accessory subunit specific to PRC2.1, which also contains PCL proteins. Unlike other accessory subunits that collectively facilitate PRC2 targeting, EPOP was implicated in an enigmatic inhibitory role, together with its interactor Elongin BC. We report an unusual molecular mechanism whereby EPOP regulates PRC2.1 by directly modulating its oligomerization state. EPOP disrupts the PRC2.1 dimer and weakens its chromatin association, likely by disabling the avidity effect conferred by the dimeric complex. Congruently, an EPOP mutant specifically defective in PRC2 binding enhances genome-wide enrichments of MTF2 and H3K27me3 in mouse epiblast-like cells. Elongin BC is largely dispensable for the EPOP-mediated inhibition of PRC2.1. EPOP defines a distinct subclass of PRC2.1, which uniquely maintains an epigenetic program by preventing the over-repression of key gene regulators along the continuum of early differentiation.

## MAIN

Polycomb repressive complex 2 (PRC2) provides a pivotal epigenetic mechanism for regulating embryonic development ^1^. A major cellular function of PRC2 in development is to catalyze the methylation of histone H3 lysine 27 (H3K27), promote the formation of repressed chromatin domains marked by trimethylated H3K27 (H3K27me3), and thereby impose epigenetic thresholds for the activation of lineage-specific genes ^2^. The gene repression activity of PRC2 is primarily mediated by at least two classes of functional PRC2 assemblies, PRC2.1 and PRC2.2, which share common core subunits and utilize distinct accessory subunits ^3^. The core subunits include the catalytic subunits and paralogs EZH1 and EZH2, EED, SUZ12, and the paralogs RBBP4 and RBBP7. The accessory subunits unique to PRC2.1 are EPOP, the homologs PALI1 and PALI2, and three mammalian homologs of *Drosophila* PCL protein—PHF1, MTF2, and PHF19, whereas those specific to PRC2.2 are AEBP2 and JARID2 ^4^. Genomic binding sites of PRC2.1 and PRC2.2 largely overlap in mouse embryonic stem cells (mESCs), suggestive of a functional redundancy ^5,6^; on the other hand, PRC2.1 and PRC2.2 can differentially impact gene expression during mESC differentiation: for example, whereas PRC2.1 preferentially maintains gene repression, PRC2.2 mainly mediates the *de novo* repression of active genes ^7^. In addition, components of PRC2.1 and PRC2.2 in mESCs display inverse chromatin binding patterns during cell cycle progression: EPOP enrichment is enhanced during the G1 phase over the S and G2/M phases, which contrasts with the case for JARID2 ^8^. Intriguingly, PRC2.1 but not PRC2.2 reduces sweet responsiveness by changing gene expression in sensory neurons in *Drosophila* ^9^.

The intricate cellular regulation of PRC2 function is primarily achieved at the level of enzymatic activity and chromatin targeting. A prominent example of the former is the allosteric stimulation of PRC2 catalysis by H3K27me3, which facilitates the spreading of the H3K27me3 repressive histone mark on chromatin through a positive feedback mechanism ^10,11^. In addition, CK2-mediated SUZ12 phosphorylation stabilizes SAM binding at the enzyme active site, promoting the ability of PRC2 in H3K27 methylation and maintenance of the differentiated cell identity ^12^. PRC2 is also directly inhibited by an oncohistone H3 and EZHIP, which contain an H3K27M or H3K27M-like protein sequence known to impede normal histone substrate binding and H3K27me3 spreading ^13–20^. The locus-specific PRC2 targeting and H3K27me3 deposition in mESCs are regulated by the accessory subunits of PRC2 ^5,6^. Particularly, AEBP2 and JARID2 bind nucleosomes with monoubiquitinated histone H2A lysine 119 (H2AK119ub1) ^21–23^, mediating PRC1-dependent chromatin recruitment of PRC2.2 ^24,25^. In comparison, PHF1, MTF2, and PHF19 link PRC2.1 to chromatin differently, via their functional domains recognizing unmethylated CpG island (CGI) DNAs ^26,27^ or trimethylated histone H3 lysine 36 (H3K36me3) active histone marks ^28–31^. Importantly, MTF2 and PHF19 can stabilize the intrinsic dimer of the PRC2 core complex, and the dimeric structure of the MTF2 or PHF19-containing PRC2.1 holocomplex enhances CGI binding, likely due to an avidity effect, which refers to the combined affinities of multivalent binding ^32^. Reminiscently, DNA binding by sequence-specific transcription factors can also be profoundly influenced by protein dimerization during active transcription ^33^.

EPOP was previously known as the PRC2 binding protein esPRC2p48 ^34^, E130012A19Rik ^35^, or C17orf96 ^36,37^. In addition to PRC2, EPOP was shown to interact with the Elongin BC heterodimer and recruit Elongin BC to PRC2 targets ^38–40^. This coincided with the reduced chromatin recruitment of PRC2 and maintenance of the low expression of some gene loci in mESCs ^38–40^. Conversely, EPOP enhanced histone methylation by PRC2 in an *in vitro* assay ^34^. In addition, EPOP appears to have PRC2- independent functions in epigenetic and transcriptional regulation at active gene loci ^38,39^. Like MTF2 and JARID2, EPOP is among the most abundant accessory subunits of PRC2 in mESCs ^34,41^, suggesting a dominant role of EPOP in gene regulation by PRC2. EPOP is extremely downregulated relative to other PRC2 subunits during neuronal differentiation ^35,41^, and it becomes highly expressed in the adult brain ^35,38,42^ (proteinatlas.org). Although important functional and structural insights on MTF2 or JARID2-containing PRC2 have started to emerge over the years ^23,32,43^, how EPOP impacts PRC2 function remains largely unclear. Here, based on a combination of biochemical, structural, and genomics data, we show that EPOP directly restricts PRC2.1 targeting to chromatin by disrupting the dimeric architecture of the enzyme complex, shaping a unique transcriptional program in mouse epiblast-like cells (EpiLCs), which mimic pregastrulating epiblasts and represent a primed state in stem cell differentiation ^44^. Our results suggest EPOP may provide an economical and efficient mechanism for regulating an epigenetic complex with a diverse array of components, preventing key developmental regulators from being over-repressed during stem cell specification.

## RESULTS

### EPOP in mESCs forms a major PRC2.1 complex displaying a distinct chromatographic behavior

During mESC differentiation, developmental genes are collectively regulated by distinct molecular assemblies of PRC2 (**Fig. 1a**). Given their high abundance, MTF2 and EPOP can theoretically assemble into at least three major PRC2.1 complexes together with the PRC2 core complex (PRC2^core^): PRC2.1^MTF2^, PRC2.1^EPOP^, and PRC2.1^MTF2–EPOP^. To assess the relative quantity of these complexes, we prepared anti-SUZ12, anti-MTF2, and anti-EPOP antibody affinity beads. We first confirmed the immunodepletion efficiency towards the respective proteins in nuclear extracts of E14 mESCs grown in 2i media (**Fig. S1**). Semi-quantitative co-immunoprecipitation (co-IP) results showed that nearly all MTF2 and about half of EPOP were bound to SUZ12 based on anti-SUZ12 co-IP, that about half of MTF2 was bound to EPOP based on anti- EPOP co-IP, and that about half of EPOP was bound by MTF2 based on anti-MTF2 co-IP (**Fig. 1b**). These results indicated that EPOP associates with a major fraction of MTF2-containing PRC2.1.

**Fig. 1:**
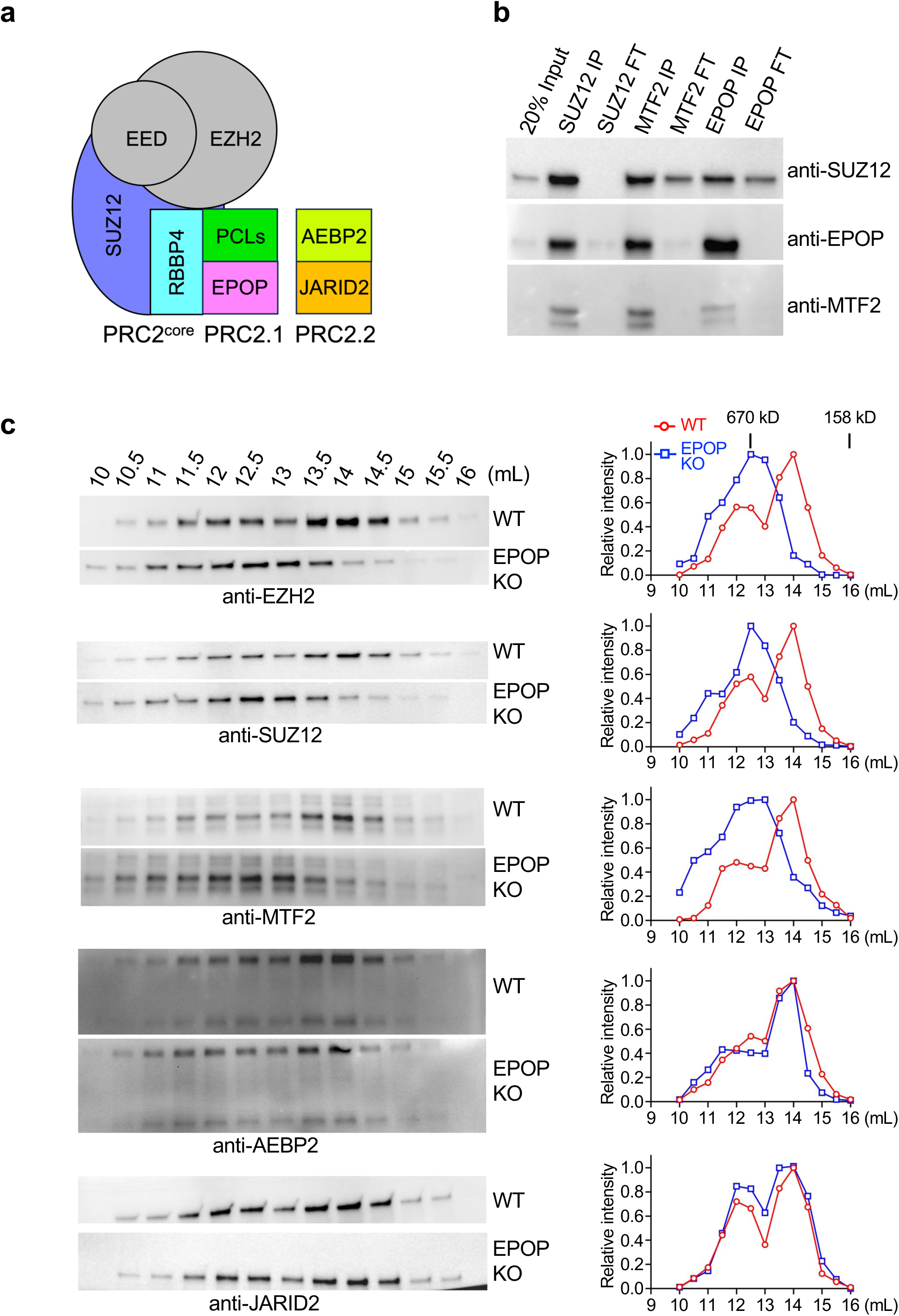
Two distinct pools of PRC2.1 in mESCs. **a**, Components of the PRC2 core complex (PRC2^core^) and the PRC2.1 and PRC2.2 holocomplexes. **b**, Co-IP assays with mESC nuclear extracts and anti-SUZ12, anti-EPOP, or anti-MTF2 beads. IP, immunoprecipitated; FT, flow through. **c**, SEC elution profiles of the nuclear extracts prepared from E14 WT or EPOP KO mESCs grown in 2i media. On the left, representative PRC2 core subunits (EZH2 and SUZ12), PRC2.1 accessory subunits (MTF2), and PRC2.2 accessory subunits (AEBP2 and JARID2) were detected by Western blot. On the right, Western blot bands were quantified and shown as the SEC elution profile.

We knocked out endogenous EPOP from E14 mESCs, which did not change the expression of other components of PRC2 (**Fig. S2**). Nuclear extracts of WT and EPOP- knockout (KO) mESCs were subjected to size-exclusion chromatography (SEC) on a Superose 6 column. MTF2 from WT mESCs was eluted as a broad peak, likely corresponding to a mixture of the monomeric and dimeric forms of MTF2-containing PRC2.1 (**Fig. 1c**) ^32^. This peak partially shifted towards earlier fractions in the absence of EPOP, suggesting accumulation of the dimeric form of PRC2.1 (**Fig. 1c**). The core subunits EZH2 and SUZ12 displayed a similar shift of the elution peak (**Fig. 1c**). In stark contrast, the PRC2.2-specific accessory subunits AEBP2 and JARID2 remained unaffected by EPOP KO (**Fig. 1c**), in line with the mutual exclusivity of PRC2.1 and PRC2.2. Together, EPOP appeared to modulate the molecular architecture of MTF2- containing PRC2.1 in mESCs.

### EPOP structurally destabilizes the PRC2.1 dimer containing MTF2 or PHF19

The role of EPOP in PRC2 remains vaguely defined, partly due to the complete absence of structural information. EPOP has been refractory to structural studies for years, given the proline-rich protein sequence and overall lack of protein structure (**Fig. S3**) ^45^. We captured the functional C-terminal domain of EPOP [EPOP(C)] bound to a PRC2.1 subcomplex in a 2.7Å crystal structure in the form of a heterotetrameric complex, which contains EPOP(C), a modified N-terminal region of SUZ12 [SUZ12(N)], RBBP4, and the reverse chromodomain of PHF19 [PHF19(RC)] ^32,38^ (**Fig. 2a and Table S1**). Importantly, the anomalous signal from the selenomethionine derivative of the L325M mutant EPOP was essential for validating the protein sequence assignment of the structural model (**Fig. 2b**). The structured portion of EPOP displays sequence homology to the relatively understudied EPOP paralog SKIDA1 and, to a lesser extent, *Drosophila* Corto, which was also shown to bridge Elongin BC to Polycomb-group (PcG) proteins ^38,40,46,47^ (**Fig. 2c**). Overall, the EPOP binding site on SUZ12 and RBBP4 is largely shared by other accessory subunits, suggesting competition in assembling distinct holocomplexes (**Fig. 2d and 2e**, also see below). In particular, we previously showed that PRC2^core^ forms a weak intrinsic dimer via domain swapping and that the dimerization interface is critically stabilized by MTF2 or PHF19 to promote chromatin targeting of PRC2.1, likely via an avidity effect ^32^. However, this dimer stabilization mechanism is disabled in the presence of EPOP (**Fig. 2d and 2e**, also see below). Key structural findings are analyzed in detail below. The resolved EPOP(C) fragment consists of four structural elements, including a short helix (SH), loop 1 (L1), a β hairpin (βH), and loop 2 (L2) (**Fig. 2b and 2f**).

**Fig. 2:**
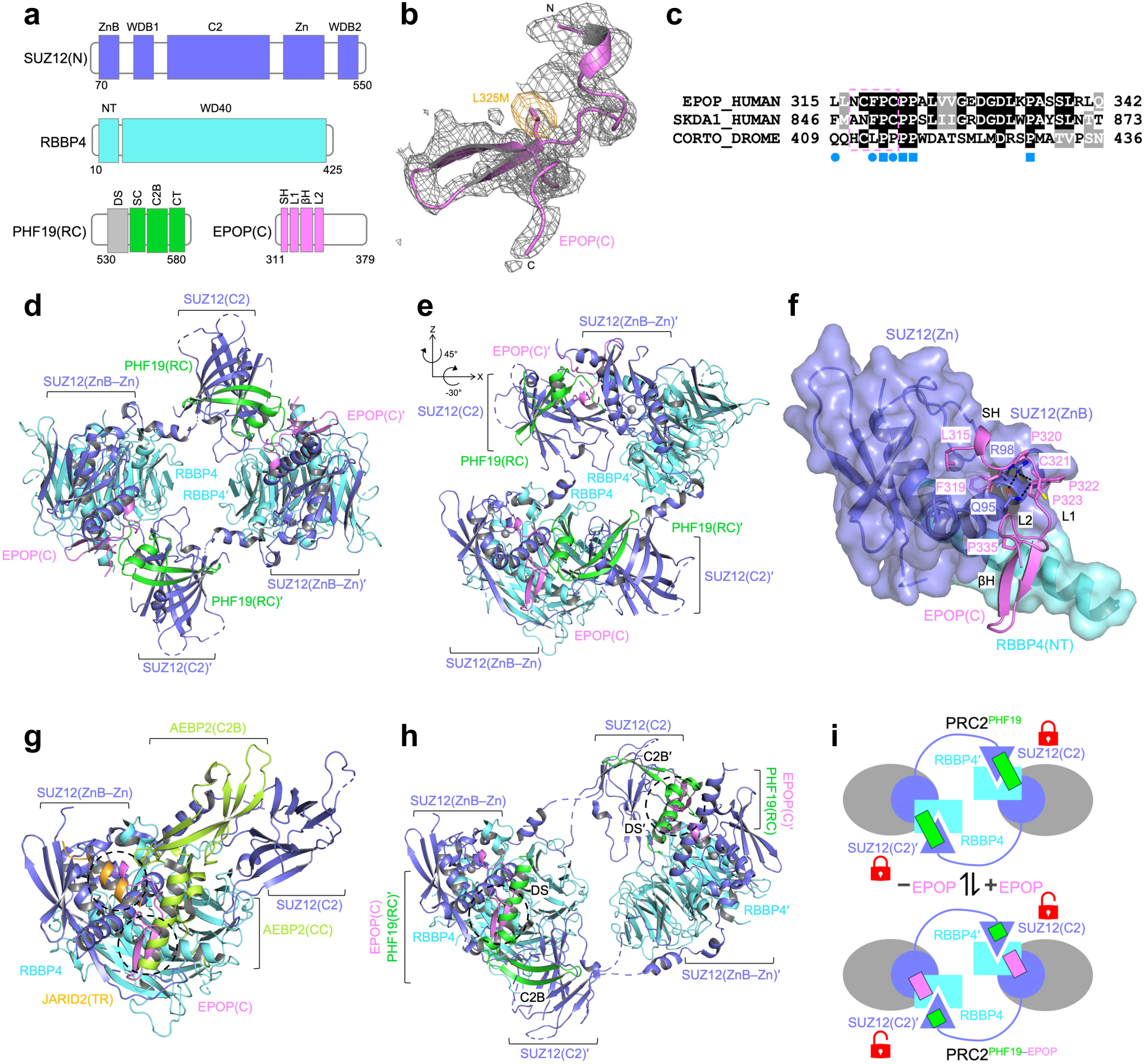
Crystal structure of an EPOP-bound PRC2.1 subcomplex. **a**, Domain structures of the proteins captured in the crystal structure. Domain names are summarized below. SUZ12(N), N-terminal domain of SUZ12, contains ZnB, zinc finger-binding helix; WDB1, WD40-binding domain 1; C2, C2 domain; Zn, zinc finger domain; WDB2, WD40-binding domain 2. RBBP4 contains NT, N-terminal domain; WD40, WD40 domain. PHF19(RC), reversed chromodomain of PHF19, contains DS, dimer stabilization helix; SC, short connecting helix; C2B, C2-binding domain; CT, C- terminal tail. EPOP(C), C-terminal domain of EPOP, contains SH, short helix; L1, loop 1; βH, β hairpin; L2, loop 2. The domain structures are color-coded based on proteins, except the DS helix of PHF19(RC), which is disordered in the structure and colored in gray. **b**, 2FoFc electron density map of the EPOP fragment contoured at 1.0σ is shown in gray. The anomalous signal of the L325M mutant contoured at 10.0σ is shown in gold. **c**, Sequence alignment of human EPOP, human SKDA1, and *Drosophila* Corto. Residues deleted in the EPOP^D5^ mutant are indicated by a dotted box. EPOP residues interacting with SUZ12 are indicated by blue discs, and proline residues contributing to the shape complementarity by blue squares. **d**, Cartoon representation of the overall structure. The SUZ12(N)–RBBP4– PHF19(RC)–EPOP(C) heterotetrameric complex adopts a dimeric structural architecture in the crystal lattice. Protein domains are labeled and color-coded. The two protomers are distinguished by the prime sign. **e**, Overall structure in a different view with the rotation matrix relative to **d** indicated. **f**, Close-up view of the EPOP binding interface. SUZ12 and RBBP4 domains are highlighted as transparent surfaces. Interacting residues are shown as sticks. Hydrogen bonds are indicated by black dotted lines. All proline residues from the EPOP fragment are also shown as sticks. **g**, Structural alignment to PRC2.2. The current structure is aligned to a PRC2.2 subcomplex containing AEBP2 and JARID2 fragments (PDBs 5WAI and 6WKR). Protein domains are labeled and color-coded. AEBP2 domains: C2B, C2-binding domain; CC, central connecting helix. JARID2 domain: TR, transrepression domain. Only the EPOP(C) fragment from the current structure is shown for clarity. **h**, Structural alignment to PRC2.1. The current structure is aligned to a PRC2.1 subcomplex containing the PHF19(RC) domain (PDB 6NQ3). Protein domains are labeled and color-coded. Only the EPOP(C) fragment from the current structure is shown for clarity **i**, Schematic model of the dimer disruption mechanism. The domain-swapped dimer is shown. The transient PRC2^core^ dimer is locked by PHF19 in PRC2.1^PHF19^. EPOP unlocks the PRC2.1^PHF19^ dimer by partially displacing PHF19.

EPOP(C)^SH^ and especially residues L315 and F319 are docked on the concave surface jointly formed by the zinc finger-binding helix and zinc finger domain of SUZ12 [SUZ12(ZnB) and SUZ12(Zn)] ^43^ (**Fig. 2c and 2f**). Proline-rich EPOP(C)^L1^ extends across the SUZ12(ZnB) helix, with residue C321 engaging in hydrogen bonding interactions with residues Q95 and R98 of SUZ12 (**Fig. 2c and 2f**). EPOP(C)^βH^ sits on the two orthogonally stacked helices, the SUZ12(ZnB) helix and the N-terminal helix of RBBP4 (RBBP4(NT)), and is secured by hydrophobic and charge interactions (**Fig. 2c and 2f**). EPOP(C)^L2^ reaches back to the SUZ12(ZnB) helix, making additional contacts (**Fig. 2c and 2f**). Notably, several proline residues of EPOP, including P320, P322, and P323 from L1, and P335 from βH, jointly contribute to the critical shape complementarity of the binding interface (**Fig. 2c and 2f**).

Structural alignments with the existing structures of PRC2.1 and PRC2.2 provided immediate insights. One concerns the assembly of PRC2.2. EPOP imposes severe steric clashes with both JARID2 and AEBP2 on the surface of the PRC2 core complex (**Fig. 2g**). Specifically, EPOP(C)^SH^ and EPOP(C)^L2^ overlap with the transrepression domain of JARID2 [JARID2(TR)], whereas EPOP(C)^L1^ and EPOP(C)^βH^ overlap with the central connecting helix of AEBP2 [AEBP2(CC)] ^32,43^ (**Fig. 2g**). In this way, EPOP may contribute to a functional balance between PRC2.1 and PRC2.2 in cells. In contrast, MTF2 or PHF19 only competes with ABEP2 but not JARID2 for PRC2^core^ binding ^32^; correspondingly, loss of AEBP2 induces the formation of a hybrid PRC2 holocomplex containing both MTF2 and JARID2 in mESCs ^3^.

We formerly showed that subdomains of the PHF19(RC) domain function as “molecular glue” to lock together the two protomers of the PRC2^core^ dimer at the domain-swapped dimer interface: whereas the C2-binding (C2B) subdomain of the PHF19(RC) domain is associated with the swapped C2 domain of SUZ12 (SUZ12(C2)) from one protomer, the dimer stabilization (DS) helix from the same PHF19(RC) molecule binds the protein body of the other protomer (**Fig. 2h and 2i)** ^32^. MTF2 likely uses the same molecular mechanism to stabilize the PRC2^core^ dimer based on the sequence homology ^32^. Remarkably, although EPOP can coexist with PHF19 in PRC2.1, the EPOP(C) fragment is incompatible with the DS helix of the PHF19(RC) domain, which becomes disordered in the current structure (**Fig. 2h and 2i**). The dimeric structural architecture of PRC2.1 is thought to facilitate chromatin targeting via an avidity effect ^32^. In this regard, EPOP is projected to impair PRC2.1^PHF19^ or PRC2.1^MTF2^ chromatin targeting by displacing the DS helix, unlocking the stabilized dimer, and thereby promoting the structural transition from the dimeric state into the monomeric state (**Fig. 2i**).

### The PRC2.1 dimer is progressively disrupted by different regions of EPOP

Inspired by the biochemical and structural observations above, we first sought to test whether EPOP can directly disrupt the PRC2.1^MTF2^ dimer in solution. All components of PRC2.1^MTF2^—EZH2, EED, SUZ12, RBBP4, and MTF2—were transiently overexpressed in HEK293T cells with or without EPOP. FLAG- and Myc-tagged EZH2 proteins were co-expressed in this context such that anti-Myc signals captured by anti-FLAG co-IP indicated the extent of PRC2.1^MTF2^ dimer formation (**Fig. 3a**). The dimeric PRC2.1^MTF2^ complex containing both FLAG-EZH2 and Myc-EZH2 was readily detected; in comparison, the presence of EPOP resulted in greatly reduced dimer formation (**Fig. 3b**), which was at least in part caused by the loss of dimer stabilization by MTF2 as predicted by the structural analysis.

**Fig. 3:**
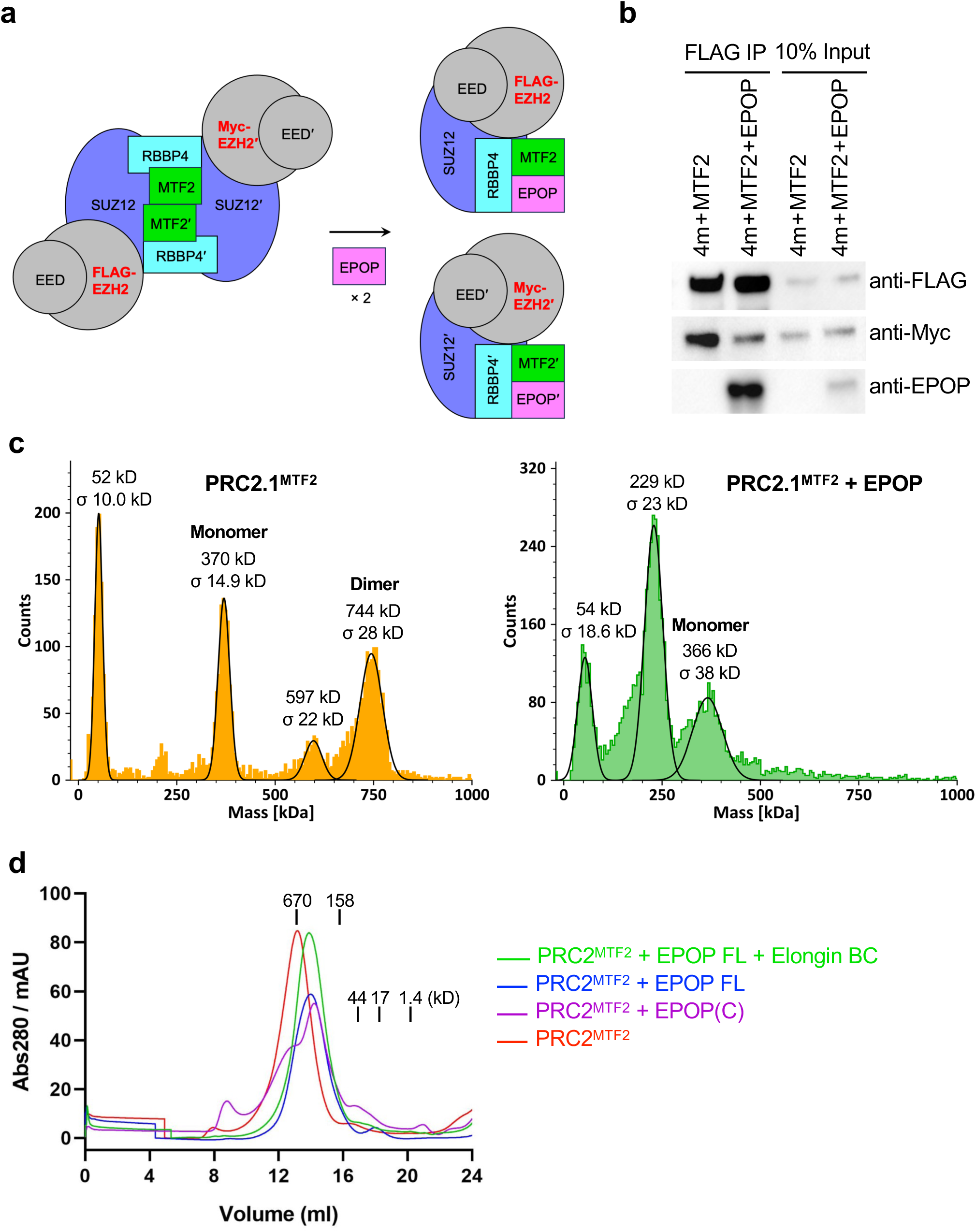
PRC2.1 dimer disruption by EPOP *in vitro*. **a**, Schematic of the co-IP-based dimer disruption assay. FLAG-EZH2 and Myc-EZH2 were co-expressed together with other indicated PRC2 subunits. **b**, Dimer disruption co-IP assay. Subunits of PRC2.1^MTF2^ were transiently expressed with or without EPOP in HEK293T cells. **c**, Mass photometry measurement. Five-member PRC2.1^MTF2^ was expressed from Sf9 cells. Full-length EPOP was added to the purified PRC2.1^MTF2^. The mass of major species in solution is indicated. Species smaller than the monomeric complex correspond to subunits or subcomplexes dissociated from the holocomplex at the 20 nM concentration. **d**, SEC elution profiles. Full-length EPOP, EPOP(C), and Elongin BC were added to the purified PRC2^MTF2^ as indicated. Protein standards are shown above the elution profiles.

Data from a reconstituted system with purified components agreed with the co-IP dimerization assay and provided additional insights. Mass photometry measurement showed that PRC2.1^MTF2^ at the concentration of 20 nM existed as an equilibrium of the monomeric and dimeric complexes and that the latter was lost upon the addition of full-length EPOP (**Fig. 3c**). We next used an orthogonal approach to assess the effect of EPOP on PRC2.1^MTF2^ dimerization. PRC2.1^MTF2^ was eluted as a dimer on a Superose 6 SEC column (**Fig. 3d**). The EPOP(C) fragment captured in the crystal structure only incompletely shifted the elution peak of the dimeric complex to a lower molecular weight (**Fig. 3d**), corresponding to a partial dimer disruption. Full-length EPOP promoted dimer disruption to a much greater extent (**Fig. 3d**), indicating EPOP regions beyond the C-terminal fragment also play a role in the dimer-to-monomer transition of PRC2.1^MTF2^.

No additional change was noticed when both EPOP and Elongin BC were supplemented into the system (**Fig. 3d**), suggesting Elongin BC, known to bind the N-terminus of EPOP, is not involved in the dimer disruption.

### Dimer disruption by EPOP weakens the chromatin binding of PRC2.1

How exactly the dimeric PRC2.1^MTF2^ complex binds chromatin is unclear. The extended homologous domain of MTF2 [MTF2(EH)] in PRC2.1^MTF2^ can mediate direct binding to the linker DNA in pairs in the context of the dimeric PRC2.1^MTF2^ holocomplex ^26,32^. A biotinylated 100 bp CGI DNA from the mouse *Lhx6* gene (CGI*^Lhx6^*) was designed to mimic the linker DNA at PRC2 binding sites and served as the bait in a pull-down assay with WT and EPOP KO mESC nuclear extracts ^32^. Bound PRC2.1 was released by restriction enzyme digestion. Noticeably more MTF2-containing PRC2.1 were captured when EPOP was absent (**Fig. 4a**), indicating EPOP restricts linker DNA binding by PRC2.1^MTF2^.

**Fig. 4:**
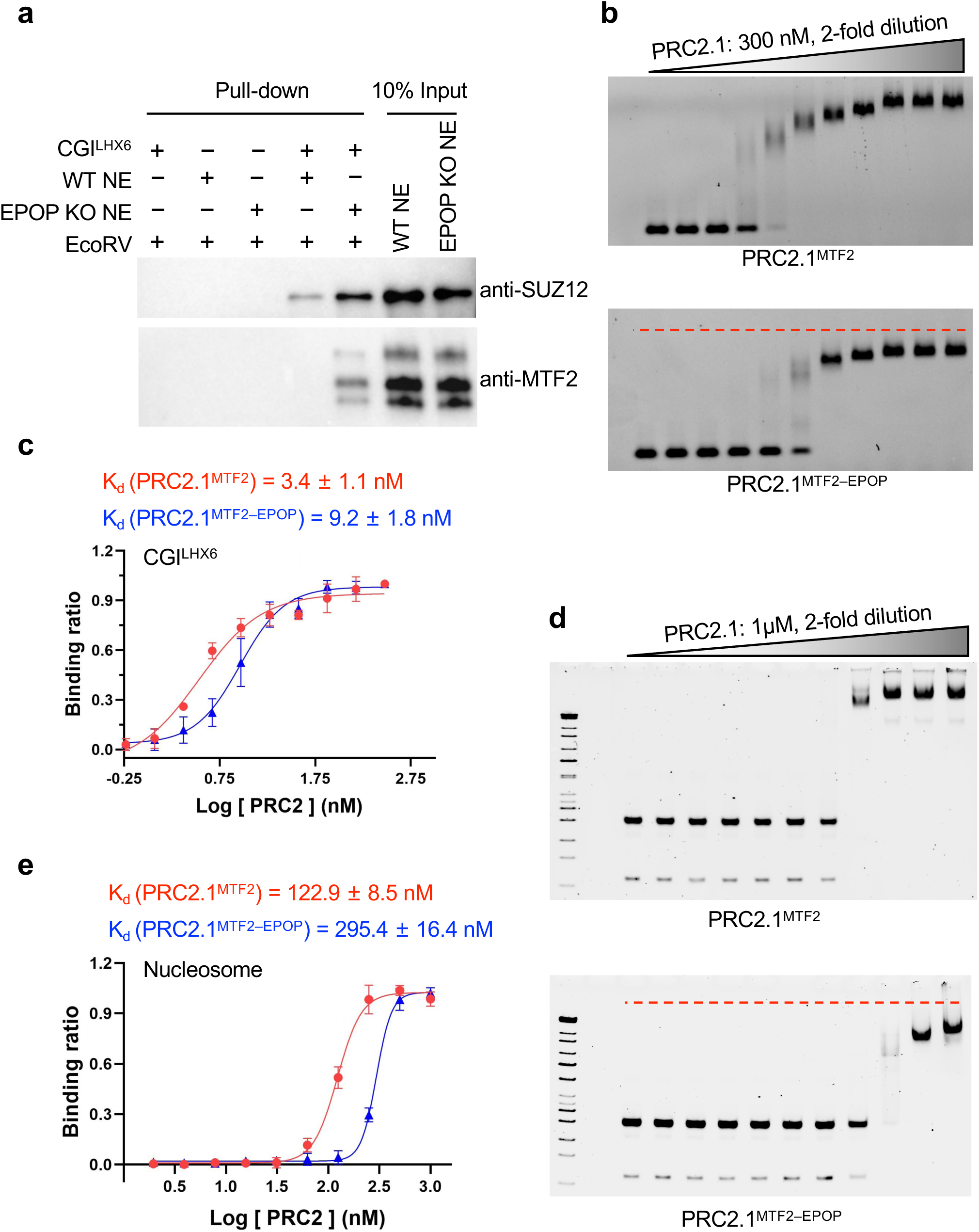
EPOP-mediated inhibition of chromatin binding by PRC2.1 via dimer disruption **a**, Nuclear extract pull-down assay. Biotinylated CGI*^Lhx6^* DNA was used as the bait. The bound PRC2.1 was released by enzyme cleavage of a restriction site inserted between biotin and the CGI*^Lhx6^*sequence. **b**, DNA binding EMSA. The serial dilution of PRC2.1^MTF2^ and PRC2.1^MTF2–EPOP^ is labeled. The dotted red line in the lower panel indicates the position of the DNA– PRC2.1^MTF2^ supercomplex to compare with that of the DNA–PRC2.1^MTF2–EPOP^ supercomplex. **c**, DNA binding affinities of the holocomplexes. **d**, Nucleosome binding EMSA. The same as **b** except that a mononucleosome was used as the probe. **e**, Nucleosome binding affinities of the holocomplexes.

To examine DNA binding quantitatively, we performed an electrophoretic mobility shift assay (EMSA) with purified PRC2.1^MTF2^ and PRC2.1^MTF2–EPOP^ (**Fig. S4**). A fluorescently labeled CGI*^Lhx6^*DNA at 0.2 nM was used as the probe, and nonspecific competitor yeast tRNA at 8 µM was added to suppress nonspecific binding.

PRC2.1^MTF2^ displayed a DNA binding affinity of 3.4 nM, and the binding affinity was reduced by 2 to 3 folds to 9.2 nM for PRC2.1^MTF2-EPOP^ (**Fig. 4b, 4c, and S5**).

Importantly, in line with the distinct oligomerization states of the enzyme complexes, DNA-bound PRC2.1^MTF2^ and PRC2.1^MTF2-EPOP^ migrated differently on the gel, with the latter behaving as a smaller supercomplex (**Fig. 4b and S5**).

Nucleosome binding was assessed by an EMSA using a ‘601’ mononucleosome with a 30 bp DNA extension on both ends. Shifted bands were clearly visible with 0.5 nM of the nucleosome probe. The measured nucleosome binding affinities of PRC2.1^MTF2^ and PRC2.1^MTF2–EPOP^ were 122.9 nM and 295.4 nM, respectively (**Fig. 4d, 4e, and S6**), indicating nucleosome binding by PRC2.1^MTF2^ is weakened by the presence of EPOP. The eight-member PRC2.1^MTF2–EPOP–Elongin^ ^BC^ complex displayed a binding affinity comparable to that of PRC2.1^MTF2–EPOP^ (**Fig. S4 and S7**), suggesting Elongin BC is unlikley involved in the EPOP-mediated inhibition of chromatin binding by PRC2.1^MTF2^. In addition, like the case for the CGI*^Lhx6^* probe, PRC2.1^MTF2^ ran as a considerably larger supercomplex when bound to the nucleosome compared to PRC2.1^MTF2–EPOP^ (**Fig. 4d and S5**), highlighting that EPOP restricts the nucleosome binding of PRC2.1^MTF2^, likely by disrupting its dimeric structure. These results suggest PRC2.1^MTF2^ and PRC2.1^MTF2–EPOP^ may display distinct chromatin binding patterns in cells, which can be further influenced by DNA sequence, epigenetic context, and transcriptional state ^48–50^.

### EPOP confers unique genome-wide profiles of PRC2.1 in EpiLCs

The enrichment of SUZ12 and H3K27me3 on chromatin was previously shown to be increased in the presence of EPOP KO or knockdown (KD) in mESCs grown in serum-containing media ^38–40^, although it remains unclear whether EPOP exerted its influence directly or indirectly. The structure of the EPOP–PRC2 complex provided a guide for a specific disruption of the complex without disturbing the PRC2-independent function of EPOP. We established isogenic mESC lines by re-expressing comparable levels of exogenous WT and mutant human EPOP proteins, EPOP^WT^ and EPOP^D5^, in the EPOP KO mESCs in 2i media (**Fig. 5a**). The EPOP^D5^ mutant containing the deletion of residues 317-321 on the binding interface displayed a severe defect in PRC2 binding (**Fig. 5b and S8**). To understand the direct role of EPOP in regulating PRC2.1 function during early development, the EPOP^WT^ and EPOP^D5^ mESCs were differentiated into EpiLCs, which mimics the cell fate transition of naïve ESCs to pregastrulating epiblasts (**Fig. 5c**) ^44^. EpiLCs also represent a transient time window for induction of the primordial germ cell-like cells (PGCLCs) *in vitro* ^44^. Unlike in further differentiated states, EPOP is still highly expressed in EpiLCs, which makes these cells a suitable model system for studying the direct PRC2-dependent role of EPOP in gene regulation.

**Fig. 5:**
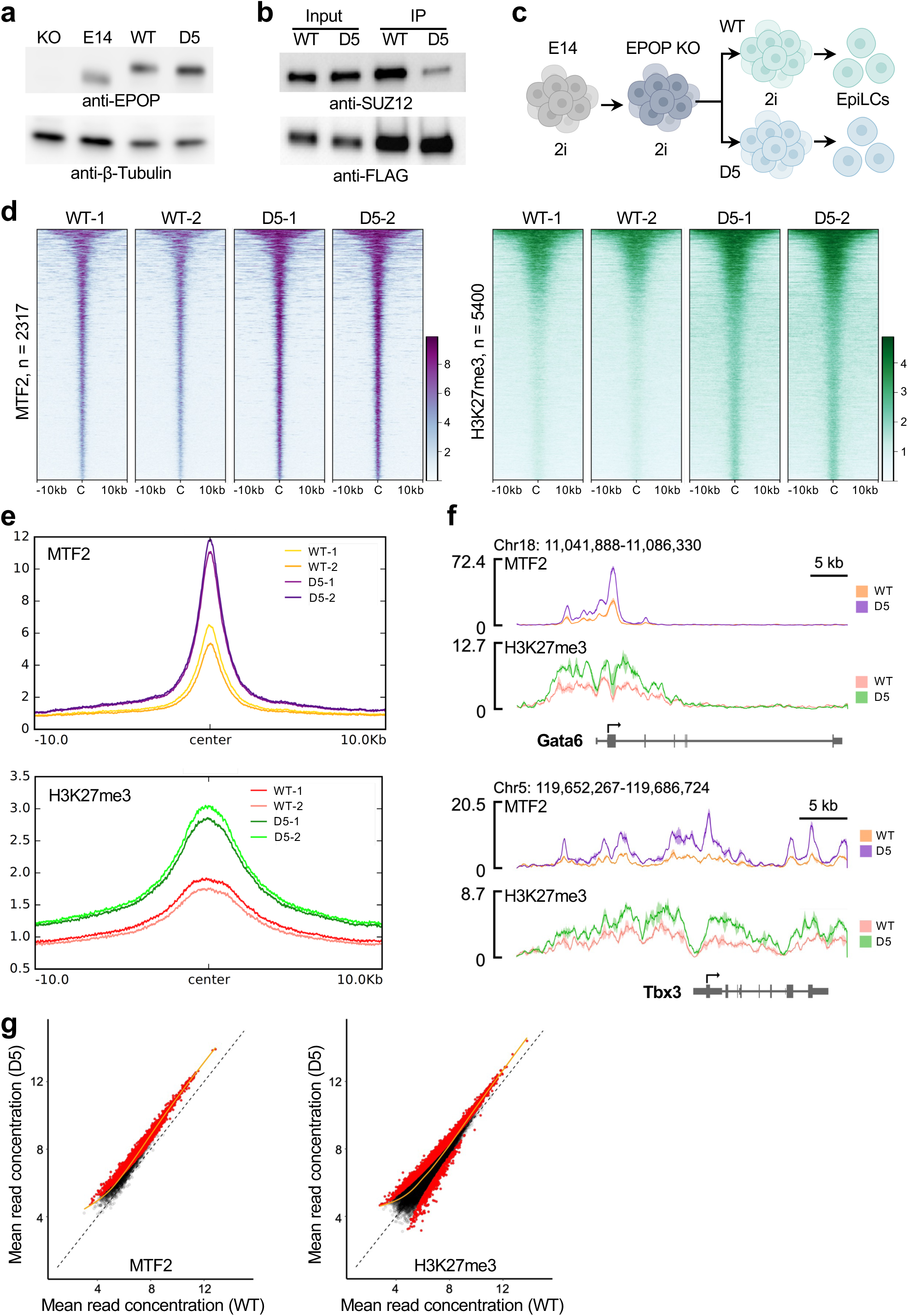
Genome-wide MTF2 and H3K27me3 enrichments regulated by EPOP in EpiLCs. **a**, Re-expression of WT and mutant EPOP in mESCs. **b**, Co-IP assay of the EPOP mutant. The re-expressed 3×FLAG-EPOP-HA protein was used as the bait, and the bound endogenous SUZ12 was detected. **c**, Schematic of the EpiLC differentiation. **d**, Heatmaps of ChIP-seq replicates. Using the FDR < 0.05 threshold, the gain-of-signal MTF2 (purple) and H3K27me3 (green) peaks from individual replicates (WT-1, WT-2, D5-1, and D5-2) are shown in the heatmaps. The centers of the consensus binding sites are aligned. The number of the consensus binding sites is labeled. **e**, Metaplots of ChIP-seq replicates. The mean ChIP-seq signal of the differential MTF2 (upper panel) and H3K27me3 (lower panel) peaks shown in **d** is plotted. The individual replicates are color-coded. **f**, Genome browser tracks of selected gene loci. Tracks were generated by SparK. Genomic coordinates are provided. The difference between the replicates calculated as the standard deviation is indicated as the shaded areas surrounding the tracks. **g**, Scatter plots of MTF2 and H3K27me3. MTF2 (left panel) and H3K27me3 (right panel) reads were normalized by the total reads. The mean read concentration corresponding to log2 (normalized ChIP-seq reads with input reads subtracted) was calculated by DiffBind.

EPOP^WT^ and EPOP^D5^ mESCs cultured in 2i media were stimulated for 48 hours to generate EpiLCs according to the established protocol ^44^. Downregulation of the naïve pluripotency genes, such as *Prdm14*, *Zfp42,* and *Esrrb,* and upregulation of EpiLC marker genes, such as *Fgf5*, *Dnmt3b*, and *Wnt3*, were observed as expected (**Fig. S9 and Table S2**). To study the specific impact of EPOP on PRC2.1, we performed chromatin immunoprecipitation followed by sequencing (ChIP-seq) to map the genome-wide distribution of MTF2 and H3K27me3. MTF2 is a dedicated accessory subunit of PRC2.1, and H3K27me3 directly correlates with the PRC2.1 function in gene repression. The consistency of ChIP-seq replicates was confirmed by the correlation plot (**Fig. S10**). Differential binding analysis of ChIP-seq peaks indicated that 2317 out of 3575 consensus binding sites showed statistically different MTF2 signals at FDR < 0.05 and that MTF2 was more enriched in the EPOP^D5^ EpiLCs than the EPOP^WT^ EpiLCs at all the 2317 sites (**Fig. 5d, 5e, and 5f and Table S3**), which is in line with the inhibitory role of EPOP in the chromatin binding of PRC2.1^MTF2^ as indicated by the *in vitro* data above. Similarly, H3K27me3 gained occupancy at 5400 out of 20742 consensus regions in the EPOP^D5^ EpiLCs compared to the EPOP^WT^ EpiLCs (**Fig. 5d, 5e, and 5f and Table S3**), indicating EPOP negatively impacts a substantial fraction of H3K27me3 loci. In contrast, 594 regions lost H3K27me3 occupancy for unknown reasons (**Table S3**). The gain of the MTF2 and H3K27me3 signals on the chromatin of the EPOP^D5^ EpiLCs was further assessed by the read count scatter plots (**Fig. 5g**). In. addition, ChIP coupled with quantitative PCR (ChIP-qPCR) was used to confirm the gain of MTF2 enrichment in two independent pairs of the EPOP^WT^ and EPOP^D5^ clones (**Fig. S11**).

### EPOP restricts PRC2.1 independently of Elongin BC

EPOP was shown to bridge the Elongin BC heterodimer to PRC2.1 ^39,40^. Elongin BC likely lost the association with PRC2.1 in the EPOP^D5^ EpiLCs. It remained ambiguous if Elongin BC contributed to the observed PRC2.1 inhibition by EPOP. To answer this question, Elongin B was stably knocked down in the EPOP^WT^ and EPOP^D5^ EpiLCs (**Fig. S12**), and the genome-wide enrichment of MTF2 was compared. In the presence of the Elongin B KD, MTF2 was differentially bound to chromatin at 3259 out of 4157 consensus binding sites, and it became more enriched at 3251—almost all the differential binding sites—in the EPOP^D5^ EpiLCs compared to the EPOP^WT^ EpiLCs (**Fig. 6a and 6b**). Notably, these sites overlapped with 85.8% of the 2813 similarly upregulated MTF2 sites in the case of the control KD (**Fig. 6a, 6b, and 6c**), indicating Elongin BC is largely dispensable for the EPOP-mediated PRC2.1 inhibition. Instead, the Elongin B KD hampered MTF2 binding at a fraction of gene loci in the EPOP^WT^ EpiLCs, which suggests Elongin BC may promote PRC2.1 targeting in EpiLCs. This phenomenon was largely lacking in the EPOP^D5^ EpiLCs (**Fig. S13**), highlighting the mediating role of EPOP in the PRC2.1–Elongin BC crosstalk. It is unclear how Elongin BC facilitates the chromatin recruitment of PRC2.1^MTF2–EPOP^ in EpiLCs; one intriguing possibility is that Elongin BC may stabilize the monomeric PRC2.1^MTF2–EPOP^ holocomplex and thereby protect it from proteasomal degradation, as shown for some other binding partners of Elongin BC ^51,52^.

**Fig. 6:**
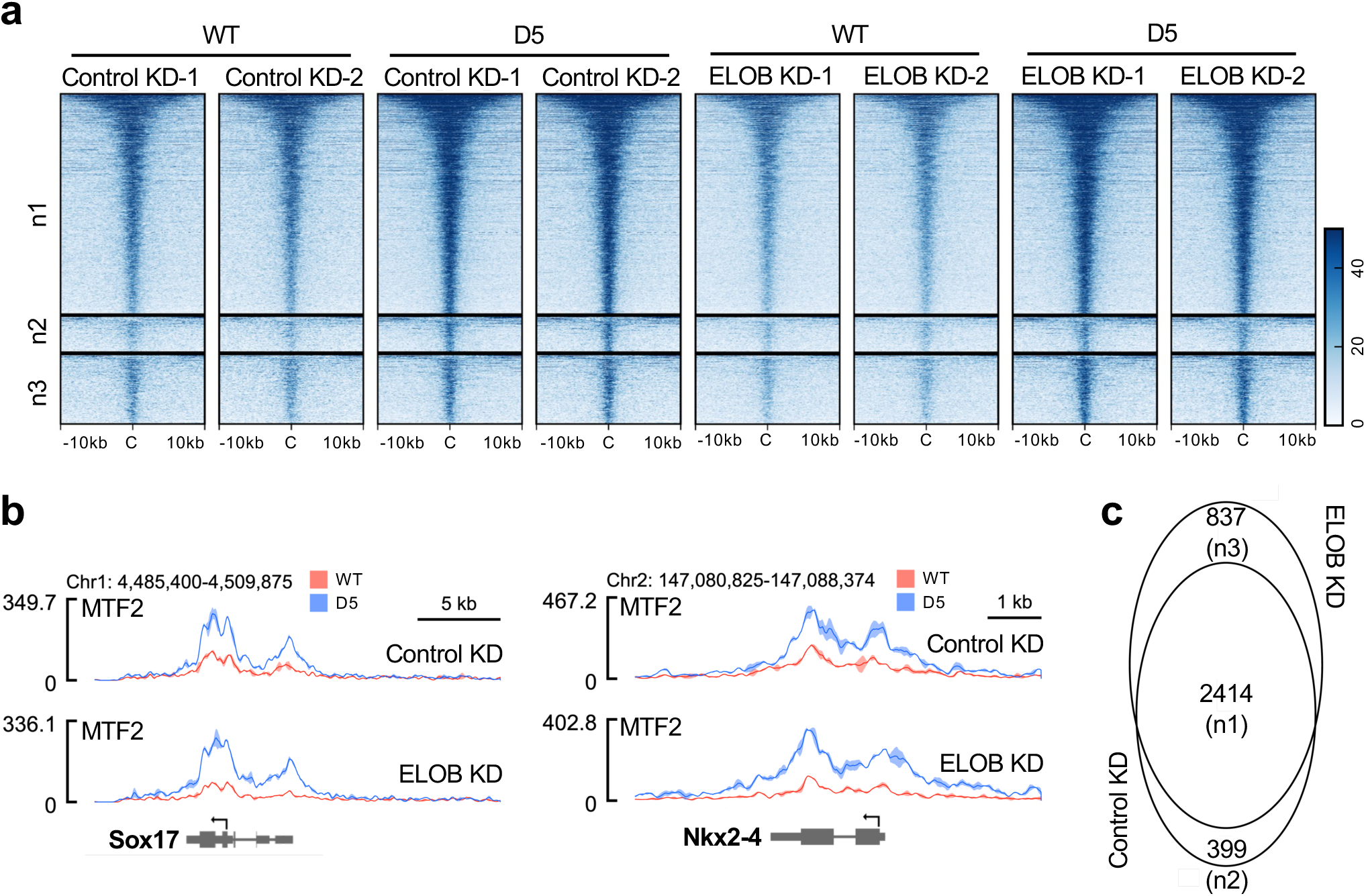
Limited role of Elongin BC in restricting PRC2.1 targeting by EPOP in EpiLCs. **a**, Heatmaps of ChIP-seq replicates. The differential MTF2 peaks between the EPOP^WT^ and EPOP^D5^ EpiLCs were grouped into three categories based on the response to shRNA KD, using the FDR<0.05 threshold. Differential peaks shared by the control and Elongin B KD are labeled as n1, differential peaks unique to the control KD are labeled as n2, and differential peaks unique to the Elongin B KD are labeled as n3. **b**, Genome browser tracks of selected gene loci. Tracks were generated by SparK. Two gene loci associated with the shared differential MTF2 peaks between the control KD and the Elongin B KD are shown. **c**, Venn diagram of the differential MTF2 peaks in the three categories.

### EPOP prevents over-repression of key developmental genes in EpiLCs

RNA-seq results revealed that the expression of a set of genes was markedly changed by the EPOP mutation in EpiLCs (**Fig. 7a and Table S2**). To directly correlate the change in gene expression with PRC2.1 targeting regulated by EPOP, the MTF2 enrichment around the transcription start site (TSS) of differentially expressed genes was compared between the EPOP^WT^ and EPOP^D5^ EpiLCs. 130 transcriptionally downregulated genes in the EPOP^D5^ EpiLCs were found to be associated with increased MTF2 signals around their TSSs, which were also accompanied by apparently higher H3K27me3 enrichments (**Fig. 7b and S14 and Table S4**). The gene ontology (GO) analysis of these PRC2.1-repressed, EPOP-maintained genes using the DAVID (Database for Annotation, Visualization, and Integrated Discovery) server indicated that sequence-specific DNA binding proteins were highly overrepresented, on the background of all the MTF2-associated genes observed (**Fig. 7c**) ^53^. These DNA binding proteins include Rel homology transcription factors, basic leucine zipper transcription factors, winged helix/forkhead transcription factors, and homeodomain transcription factors, which are associated with biological processes like regulation of transcription, stem cell maintenance, and organ development (**Fig. S15**).

**Fig. 7:**
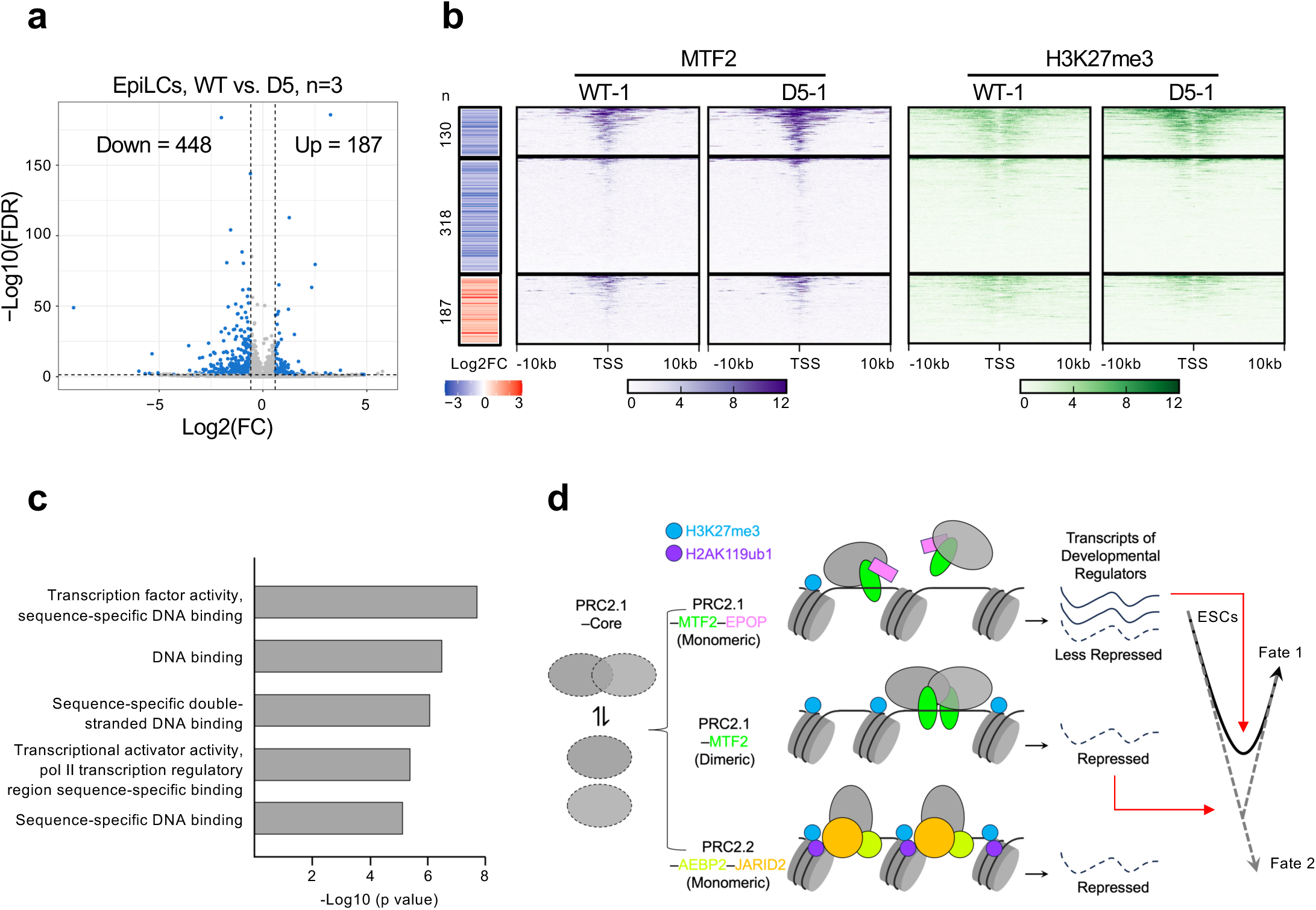
Developmental gene regulation by EPOP through PRC2.1. **a**, Volcano plot of differential gene expression. RNA-seq results of the EPOP^D5^ EpiLCs in triplicates were compared to those of the EPOP^WT^ EpiLCs. The number of upregulated and downregulated genes is indicated. Data passing the FDR < 0.05, FC > 1.5, and average TPM of WT or mutant > 0.5 thresholds were analyzed. **b**, Correlation of RNA-seq and ChIP-seq. One ChIP-seq replicate is shown here, and the other replicate is shown in the supplemental materials. The differential gene expression was aligned with the differential MTF2 enrichment around the transcription start site (TSS). The corresponding H3K27me3 signals are also displayed. Compared to the WT counterpart, 130 genes in the EPOP^D5^ EpiLCs were downregulated and associated with enhanced MTF2 signals around the TSS. Other 318 downregulated genes were not associated with EPOP-regulated MTF2 targeting. 187 upregulated genes are shown as well. **c**, Gene ontology analysis. The PRC2.1-repressed, EPOP-maintained genes were subjected to gene ontology analysis on the DAVID server. The top 5 overrepresented terms in molecular function are shown. **d**, Schematic model of developmental gene repression by PRC2. On the left, the transient intrinsic dimer of the PRC2 core complex is illustrated. In the middle, distinct oligomerization states of various PRC2.1 and PRC2.2 holocomplexes are highlighted. MTF2 mediates direct chromatin binding, and it also stabilizes the intrinsic dimer, promoting chromatin targeting of the dimeric PRC2.1, likely via an avidity effect. EPOP disrupts the dimeric architecture of PRC2.1 containing MTF2, restricts PRC2.1 targeting, and thereby maintains the limited expression of PRC2.1-repressed developmental regulators. On the right, the PRC2.1-dependent role of EPOP in early development is illustrated on the right. Black solid curve: during the ESC differentiation, a set of key gene regulators is repressed by PRC2.1, with limited expression being maintained by the EPOP-mediated inhibition of PRC2.1 targeting, which is followed by upregulation of the same set of gene regulators, leading to cell fate 1. Gray dotted curve: the absence of EPOP results in the over-repression of these gene regulators by PRC2.1, which may change stem cell differentiation trajectories.

Although targeted for repression during the EpiLC differentiation, some of the PRC2.1-repressed, EPOP-maintained genes may still play a crucial role in the next step of cell fate specification. For example, in agreement with the prediction, five of these genes, Esrrb, Gjb3, Plagl1, Prdm1, and Prdm14, were found in the list of top 100 upregulated genes during the EpiLC-to-PGCLC transition, and none in the downregulated gene list (**Table S4**) ^44^. Therefore, by directly disrupting the dimeric form of PRC2.1 and thereby restricting its chromatin targeting, EPOP appears to act as a “brake” on the repressive system to prevent abrupt over-repression of key developmental regulators during the epiblast differentiation, which might conceivably help establish the critical time window for the downstream PGC specification (**Fig. 7d**)^44^.

## DISCUSSION

PRC2.1 and PRC2.2 are the two distinct classes of the PRC2 holocomplexes. Emerging evidence indicates that PRC2.1 and PRC2.2 are involved in non-overlapping processes in cell development, which is largely dictated by their respective accessory subunits. Although these accessory subunits are believed to collectively contribute to the chromatin recruitment of PRC2 and H3K27me3 deposition ^5,6^, EPOP, an accessory subunit of PRC2.1, is an exception. EPOP was formerly noted to somehow limit PRC2 binding to chromatin ^39,40^, and the present work provides a molecular mechanism for this puzzling observation. Our previous work showed that PRC2.1 containing only MTF2 or PHF19 forms a domain-swapped dimer, which facilitates chromatin targeting of the holocomplex likely via an avidity effect ^32^. Here, we find EPOP can directly disrupt the dimeric form of PRC2.1 and thereby constrains its chromatin binding both *in vitro* and *in vivo*. Both transcription factors and epigenetic complexes carry out their functions by interacting with chromatin; whereas the role of transcription factor dimerization in transcription has been well recognized ^33^, the present study exemplifies how the oligomerization state of a central epigenetic complex can be regulated to impact biological output during development (**Fig. 7d**).

The crystal structure of an EPOP-bound PRC2.1 subcomplex lays an important foundation for understanding the mechanistic model of the dimer disruption. Anomalous signals are essential for removing some ambiguities in the structural model of the C-terminal proline-rich region of EPOP. This EPOP region disables the mechanism of the MTF2 or PHF19-mediated dimer stabilization of PRC2.1 by displacing the dimer stabilization helix of MTF2 or PHF19 from the core complex, which allows other EPOP regions to complete the dimer disruption process. Despite the partial structural displacement of MTF2 and PHF19 by EPOP, they can still be stably and stoichiometrically associated with the PRC2 core complex. How the largely unstructured EPOP region beyond the PRC2-interacting C-terminal region contributes to the dimer disruption is unknown. In contrast, PHF1 lacks the structural element responsible for dimer stabilization ^32^, and it remains to be determined if PHF1-containing PRC2.1 is similarly regulated by EPOP in cells.

In the reconstituted system, the direct consequence of PRC2.1^MTF2^ dimer disruption by EPOP is the substantially weakened binding of linker DNA and nucleosome *in vitro*, which is also largely recapitulated by ChIP-seq *in vivo* in EpiLCs. The structure-based EPOP mutant defective in PRC2 binding brings invaluable specificity to the *in vivo* study, particularly considering EPOP may also function in a non- PRC2 context ^39^. The dimeric PRC2.1^MTF2^ and monomeric PRC2.1^MTF2–EPOP^ represent two major pools of the PRC2.1 holocomplex in mESCs. In the EpiLCs, the loss of the EPOP–PRC2.1 interaction converts PRC2.1^MTF2–EPOP^ into PRC2.1^MTF2^, resulting in the genome-wide increase of the MTF2 enrichment, likely due to the enhanced chromatin binding of the dimeric complex as compared to the monomeric complex. Accordingly, the genomics data may now be understood mechanistically on the molecular level.

The combination of the ChIP-seq and RNA-seq data reveals a list of PRC2.1- repressed, EPOP-maintained genes in EpiLCs, which become over-repressed when EPOP no longer binds PRC2.1. Sequence-specific transcription factors are overrepresented on the background of all the detected MTF2-associated genes, indicating the EPOP-mediated inhibition of PRC2.1 maintains the partial expression of a group of key gene regulators. These genes start to be repressed by PRC2.1 during the ESC-to-EpiLC differentiation but may remain needed for downstream developmental processes. Indeed, some of these genes are known to be activated during the EpiLC-to-PGCLC cell fate transition ^44^. In this regard, we speculate that EPOP may impact early development by tempering the sharp change in the gene expression profile (**Fig. 7d**). Apparently, compared to the coordinated transcriptional regulation of multiple subunits of PRC2.1, the single EPOP protein provides an economical mechanism to modulate the PRC2.1 activity in cells in a specific and efficient way. Although we are focused on EpiLCs here, it is interesting to posit that EPOP may similarly regulate gene expression in the nervous system, where this protein is also found to be expressed.

Elongin BC is an interactor of EPOP ^39,40^, and the role of Elongin BC in gene regulation by PRC2 remains vaguely defined. Our data suggest that Elongin BC is not involved in the PRC2.1 dimer disruption by EPOP and is largely dispensable for the EPOP-mediated inhibition of PRC2.1 targeting both *in vitro* and in EpiLCs. Interestingly, Elongin BC appears to promote MTF2 binding to chromatin at a set of gene loci in an EPOP-dependent manner. Although experimental evidence is lacking, one possibility is that Elongin BC may help stabilize the monomeric PRC2.1^MTF2–EPOP^ in cells, which may become less stable than PRC2.1^MTF2^ due to the dimer disruption. Similarly, Elongin BC was previously shown to stabilize the SOCS1 suppressor of cytokine signaling protein and VHL tumor suppressor protein by forming a protein complex and protecting the respective protein from proteasomal degradation ^51,52^.

## DATA AVAILABILITY

The crystal structure solved in this work has been deposited in the Protein Data Bank under the accession number 9B32. Genomics data have been deposited in the Gene Expression Omnibus under the accession numbers GSE264672 and GSE264673.

## ACKNOWLEDGMENTS

The cDNA of EPOP was kindly provided by Dr. Robert Liefke and Dr. Yang Shi. The cDNAs of human PRC2 core subunits were kindly provided by Dr. Robert Kingston. This research was supported by NIH grant R35GM136308 to X.L. X.L. is a W.W. Caruth, Jr., Scholar in Biomedical Research. L.G. was supported by American Heart Association postdoctoral fellowship 19POST34450043. S.C. was supported by National Natural Science Foundation of China grant 32100464. Some data presented in this report were acquired with a mass photometer that was supported by award S10OD030312-01 from the NIH. Results shown in this report are derived from work performed at Argonne National Laboratory, Structural Biology Center (SBC) at the Advanced Photon Source. SBC-CAT is operated by UChicago Argonne, LLC, for the U.S. Department of Energy, Office of Biological and Environmental Research under contract DE-AC02-06CH11357. Use of the Stanford Synchrotron Radiation Lightsource, SLAC National Accelerator Laboratory, is supported by the U.S. Department of Energy, Office of Science, Office of Basic Energy Sciences under Contract No. DE-AC02-76SF00515. The SSRL Structural Molecular Biology Program is supported by the DOE Office of Biological and Environmental Research, and by the National Institutes of Health, National Institute of General Medical Sciences (P30GM133894). The contents of this publication are solely the responsibility of the authors and do not necessarily represent the official views of NIGMS or NIH.

## AUTHOR CONTRIBUTIONS

Xin.L. conceived the study; L.G., Xiuli.L., X.Y., S.C., and Xin.L. designed and performed the experiments; X.Y. and Xin.L. analyzed the structural data; Xiuli.L., Z.Y., C.X., and Xin.L. analyzed the genomics data; L.G., Xiuli.L., X.Y., Z.Y., C.X., and Xin.L. wrote the manuscript.

## METHODS

### Antibodies

Antibodies used in this study are listed below: SUZ12 antibody (CST, Cat. # 3737S), MTF2 antibody (Proteintech, Cat. # 162081AP), EPOP antibody (Active Motif, Cat. # 61753), JARID2 antibody (CST, Cat. # 13594S), AEBP2 antibody (CST, 14129S), Elongin B antibody (Abcam, Cat. # ab168836), EZH2 antibody (CST, Cat. # 5246S), FLAG antibody (Sigma, Cat. # F1804), Myc antibody (CST, Cat. # 2276S), H3 antibody (CST, Cat. # 9715S), H3K27me3 antibody (CST, Cat. # 9733S), β-Tubulin antibody (CST, Cat. # 2128S).

### mESC culture

The parental E14 mESCs (ES-E14TG2a) were obtained from ATCC. Cells were cultured on 0.2% gelatin-coated plates in 2i/LIF medium (1:1 mix of DMEM/F-12 and Neurobasal medium supplemented with 1 × Pen/Strep, 0.05% BSA, 1 mM GlutaMAX, 100 μM β-mercaptoethanol, 0.5 × N2 supplement, 1 × B27 supplement, 1 μM MEK inhibitor PD0325901, 3 μM GSK3 inhibitor CHIR99021, and home-made LIF). Medium was changed each day and cells were passaged every 2 or 3 days. Cells were tested routinely for mycoplasma (Bulldog Bio Inc.) e-Myco PLUS Mycoplasma PCR Detection Kit (Bulldog Bio Inc.)

### EpiLC differentiation

EpiLC induction was performed according to a previous publication ^44^. Briefly, 1.5 × 10^6^ mESCs were plated on a 10 cm dish coated with human fibronectin (16.7 μg/ml) in N2B27 medium (1:1 mix of DMEM/F-12 and Neurobasal medium supplemented with containing 1 × Pen/Strep, 2 mM GlutaMAX, 50 μM β-mercaptoethanol, 0.5 × N2 supplement, 0.5 × B27 supplement, 20 ng/ml activin A, 12 ng/ml bFGF, and 1% KSR). The medium was replaced every day until 48 hours before cell collection. The differentiation was confirmed using RT-qPCR for pluripotency and EpiLC-specific markers.

### Establishment of EPOP KO mESC lines

EPOP KO mESC lines were generated by transfecting E14 mESCs with the pSpCas9 (BB)-2A-GFP (PX458) vector (Addgene) containing the gRNA targeting EPOP Exon 1 (CGAGCAGGGAGACCCCCGCG) ^5^. Electroporation was performed using the Lonza transfection system. GFP-positive cells were sorted on day 3 and plated on 96-well plates after dilution of the sorted cells for single-cell clone selection. Western blot and Sanger sequencing were used to confirm the EPOP KO.

### Re-expression of EPOP^WT^ and EPOP^D5^

WT and D5 mutant EPOP proteins were re-expressed in EPOP KO mESCs using the pCDH-EF1a-3×FLAG-EPOP-HA construct with a blasticidin selection marker. The Kozak sequence was inserted to improve the expression. HEK293T cells were transfected with the EPOP^WT^ or EPOP^D5^ lentiviral constructs, together with psPAX2 and Vsvg plasmids using Xtreme Gene 9 DNA transfection reagent (Roche). The culture medium containing the lentivirus was collected and concentrated using a Lenti-X^TM^ concentrator (Takara) overnight. Concentrated lentivirus was added to EPOP KO mESCs in the presence of polybrene (8 ug/ml) overnight. 48 hours post-transfection, cells were selected in the presence of 10 μg/ml blasticidin for 7 days. Single-cell clones were generated in 96-well plates. Cells were collected for Western blot analysis to pick clones expressing comparable EPOP^WT^ and EPOP^D5^.

### shRNA KD

The shRNA targeting the Elongin B gene TCEB2 (GATGTGATGAAGCCACAGGAT) was purchased from Sigma. For TCEB2 KD, the lentivirus was produced as described above. EPOP^WT^ and EPOP^D5^ mESCs were transfected with the concentrated lentivirus in the presence of polybrene (8 ug/ml). 48 hours post-transfection, puromycin (1 μg/ml) was added to the medium for selection. After 7 days of selection, cells were collected for Western blot analysis.

### ChIP-qPCR ChIP-Seq

Cells were trypsinized with TrypLE™ Express Enzyme and washed once with PBS before crosslinking with 1% formaldehyde for 10 minutes. Formaldehyde was quenched with 0.125 M Glycine for 5 minutes before two PBS washes. The crosslinked cells were lysed in Farnham lysis buffer (5 mM PIPES pH 8.0, 85 mM KCl, 0.5% NP40, 1 mM DTT, and 1× protease inhibitor cocktail) to collect nuclei. Nuclei were resuspended with the lysis buffer (50 mM Tris-HCl, pH 7.9, 10 mM EDTA, 1% SDS, 1 mM DTT, and 1× protease inhibitor cocktail) and sonicated (Diagenode Bioruptor) using 30 s ON/30 s OFF for 4 cycles. Insoluble chromatin was pelleted by centrifugation at 15,000 g at 4 °C for 10min. The supernatant was harvested to determine the chromatin concentration and sonication efficiency. The sheared chromatin was diluted 10-fold with ChIP dilution buffer (20 mM Tris-HCl, pH 7.9, 2 mM EDTA, 150 mM NaCl, 0.5% Triton X-100, 1 mM DTT, and 1× protease inhibitor cocktail) for ChIP. 50 μg chromatin was used for MTF2 ChIP and 20 μg for H3K27me3 ChIP. Chromatin was incubated with 5 μg antibodies while rotating at 4 °C overnight. The next day, the antibody-chromatin complexes were incubated for 2 hours at 4 °C with Protein A/G Dynabeads. After incubation, the beads were washed with 1 mL of each of the following wash buffers in this order: low salt buffer (20 mM Tris-HCl, pH 8.0, 2 mM EDTA, 1% Triton X-100, 0.1% SDS, and 150 mM NaCl), high salt buffer (20 mM Tris-HCl, pH 8.0, 2 mM EDTA, 1% Triton X-100, 0.1% SDS, and 500 mM NaCl), LiCl buffer (10 mM Tris-HCl, pH 8.0, 1 mM EDTA, 1% NP40, 1% sodium deoxycholate, and 250 mM LiCl), and TE buffer (20 mM Tris-HCl, pH8.0, and 1 mM EDTA). Samples were eluted with the elution buffer (1% SDS and 100 mM NaHCO3) followed by reverse crosslinking overnight. RNase treatment and Proteinase K digestion were performed before phenol: chloroform: isoamyl alcohol (25:24:1) and chloroform extraction.

For deep sequencing, 8 ng of eluted DNA was used to build the library with the NEBNext Ultra II kit (NEB) according to the manufacturer’s protocol, except that Ampure XP beads were used for the size selection. Libraries were sequenced on Illumina NextSeq 2000 (UT Southwestern) or Novaseq X plus (Novogene) using the paired-end strategy. For quantification by qPCR, LightCycler 480 Instrument II system and SYBR^TM^ Green PCR master mix were used. 3 μl of diluted ChIP DNA was used for each PCR reaction. Statistical analysis was performed using Microsoft Excel and GraphPad Prism. Primer sequences are provided below.

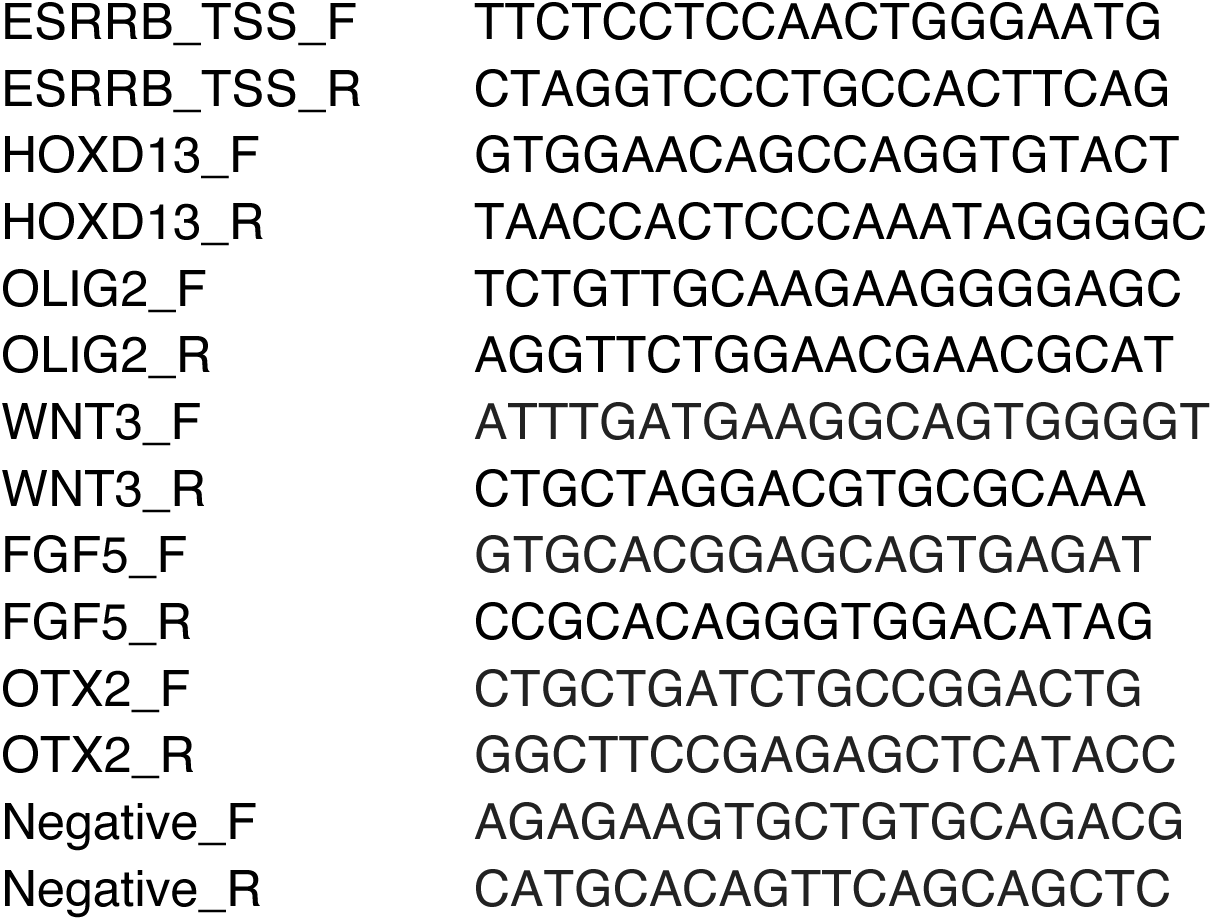

### RNA-Seq

Total RNA was extracted using TRIzol^TM^ reagent (Invitrogen) according to the manufacturer’s instructions. The concentration and purity of RNA were assessed using a NanoDrop device (Life Technologies). RNA integrity was checked on a TapeStation system (Agilent Technologies). The RNA-seq library was prepared using NEBNext Ultra™ II RNA Library Prep Kit for Illumina (NEB), and deep sequencing was conducted using the PE150 strategy on NovaSeq 6000 platform.

### Bioinformatics analysis

For the initial ChIP-seq data without the shRNA KD sequenced at UT Southwestern, the paired-end 40 bp read length FASTQ files were checked for quality using FastQC and FastQ Screen. Reads were aligned to the mm10 reference genome with Bowtie 2 ^54^, and alignments with mapping quality scores less than 10 were filtered out. Duplicates were removed using SAMtools ^55^. ChIP-seq peaks were called using input ChIP samples as controls ^56^. BAM coverage BigWig files were generated by bamCoverage in deepTools ^57^. The differential binding analysis was performed using DiffBind ^58^, with normalization by total reads and blacklist regions removed. *Drosophila* spike-in reads were present but not used for normalization due to large variations. For the shRNA KD ChIP-seq data sequenced by Novogene, the raw sequencing reads were trimmed by Cutadapt and then mapped by Bowtie 2 with the “--very-sensitive” parameter. The aligned reads were filtered by SAMtools using the “-F 1804 -f 2 -q 30” parameter.

Duplicates were removed by Picard. ChIP-seq peaks were called by MACS2 using the “-f BAMPE -q 0.05 --broad” parameter. Differential peaks were identified by DiffBind using FDR<0.05 as the cutoff.

The read densities ± 10 kb surrounding the peak center or TSS were generated and visualized as heatmaps using deepTools. Metagene analyses were performed to show the distribution of average ChIP-seq read densities ± 10 kb surrounding the peak center or TSS using deepTools. Genome browser tracks with the standard deviation of replicates were plotted with Spark (https://github.com/harbourlab/SparK), an NGS visualization tool. The mean read concentration of each condition calculated by DESeq2 was applied to generate scatter plots using the R package ggplot2. The Venn diagram was generated using the R package VennDiagram. For RNA-seq, differentially expressed genes between the EPOP^WT^ and EPOP^D5^ EpiLCs were associated with differential MTF2 binding around TSSs. The expression fold changes were used to plot the heatmap using the R package ggplot2. GSEA was performed with GSEA software (https://www.gsea-msigdb.org/gsea/downloads.jsp). Pre-ranked gene lists were based on differential expression analysis. Genes activated during the EpiLC differentiation were taken from a previous study and defined as the EpiLC-specific genes ^59^. R package ggplot2 was used to generate the volcano plot and the MA plot. The deepTools suite was used to generate the Pearson correlation heatmap. The read coverage statistics for genomic regions for each BAM file are calculated by multiBamSummary, and the correlation matrix using Pearson’s correlation method is plotted with plotCorrelation.

### mESC nuclear extract preparation

WT and EPOP KO E14 mESCs were harvested at around 70-80% confluence from 10 cm dishes and washed twice with cold PBS. 5 volumes of hypotonic buffer A (10 mM HEPES, pH 7.9, 10 mM KCl, 1.5 mM MgCl2, 0.5% NP40, 1mM DTT, 1mM PMSF, and 1× protease inhibitors) were added to the pellet to lyse the cells. Nuclei were collected by centrifugation at 1000× g at 4°C for 10 min. 5 volumes of ice-cold high-salt buffer (20 mM HEPES, pH 7.9, 420 mM KCl, 1.5 mM MgCl2, 0.2 mM EDTA, 25% glycerol, 1mM PMSF, and 1× protease inhibitors) were added to lyse the nuclei. The mixture was incubated on a rotator at 4°C for 1 hour, followed by centrifugation at 15,000× g at 4°C for 10 min. Nuclear extracts in the supernatant were kept.

### Size exclusion chromatography

For the SEC of nuclear extracts, nuclear extracts were dialyzed against the SEC buffer (20 mM Tris-HCl, pH 8.0, 150 mM NaCl, and 2 mM DTT), followed by centrifugation at 15,000× g at 4°C for 10 min. The supernatant containing 200 μg nuclear extracts was loaded onto an Superose 6 10/300 GL column (GE Healthcare) in the SEC buffer.

Fractions were collected and analyzed with Western blot. For the SEC of purified PRC2.1 complexes, PRC2.1^MTF2^ was mixed with EPOP (residues 311-379), full-length EPOP-1-379, or full-length EPOP plus Elongin BC at the 1:5, 1:5, or 1:5:5 molar ratio, respectively. The mixed proteins were dialyzed against the reconstitution buffer (20 mM Tris-HCl, pH 8.0, 300 mM NaCl, 10% glycerol, and 2 mM DTT, at 4°C overnight.

Reconstituted complexes were loaded onto an Superose 6 10/300 GL column in the SEC buffer (20 mM Tris-HCl, pH 8.0, 300 mM NaCl, and 2 mM DTT), and the elution profile was recorded.

### Expression and purification of PRC2.1^MTF2^, PRC2.1^MTF2–EPOP^, and PRC2.1 ^MTF2–EPOP– Elongin BC^

PRC2.1^MTF2^ (EZH2–EED–SUZ12–RBBP4–MTF2) were expressed in Sf9 insect cells as described previously ^32^. Cell pellets were harvested, resuspended in the lysis buffer (50 mM Tris-HCl, pH 8.0, 1 M NaCl, 1 mM PMSF, 1× protease inhibitors, 0.1% NP40, 10% glycerol, and 2 mM DTT), and sonicated. The cell lysate was clarified, and proteins were captured using the IgG affinity resin. The bound proteins were released by the TEV cleavage overnight at 4°C. The eluted proteins were dialyzed against the ion exchange low salt buffer (50 mM sodium citrate, pH 6.0, 50 mM NaCl, and 2 mM DTT) and further purified by the Mono S 5/50 GL column, with a gradient wash from the ion exchange low buffer to the ion exchange high salt buffer (50 mM sodium citrate, pH 6.0, 1 M NaCl, and 2 mM DTT). PRC2.1^MTF2^ was finally purified on a Superose 6 10/300 GL column in the SEC buffer (20 mM Tris-HCl, pH 8.0, 300 mM NaCl, and 2 mM DTT). PRC2.1^MTF2–EPOP^ was expressed and purified similarly. PRC2.1^MTF2–EPOP^ was mixed with bacterially expressed Elongin BC heterodimer to reconstitute PRC2.1^MTF2–EPOP–Elongin^ ^BC^.

### Mass photometry

200 nM PRC2.1^MTF2^ was incubated with or without 600 nM full-length EPOP in the fresh MP buffer (20 mM Tris-HCl, pH 8.0, 150 mM NaCl, and 1mM DTT) on ice for 30 min.

Sample chambers were assembled with a 2 × 4 well gasket on the clean cover slide and were placed on the Two MP mass photometer (Refeyn). 16.2 μl MP buffer was loaded onto one sample well and focused, followed by adding a 1.8 μl sample and mixing in the well. The AcquireMP software package was used to record contrast dots for 1 minute, and the data were analyzed with the DiscoverMP software.

### Co-IP of PRC2.1 complexes in nuclear extracts

The antibody-conjugated resin (anti-MTF2, anti-EPOP, or anti-SUZ12) was made using cyanogen bromide-activated-Sepharose 4B (Sigma) following the manufacturer’s instructions. Antigen binding efficiency was checked using the immunodepletion method. Nuclear extracts of WT or EPOP KO E14 mESCs were mixed with the homemade antibody resin and washed with the wash buffer (50 mM Tris-HCl, pH 8.0, 150 mM NaCl, 0.1% NP40, 2mM DTT, and 10% Glycerol). Captured protein complexes were released with 200 mM glycine, pH 2.8, and analyzed using Western blot.

### Co-IP dimerization assay

pCS2-Myc-EZH2, pCS2-FLAG-EZH2, pCS2-His6-EED, pCS2-HA-SUZ12, pCS2-HA-RBBP4, pCS2-ProteinA-3C-MTF2, and pCS2-ProteinA-TEV-EPOP plasmids were transiently co-transfected into HEK293T cells to express PRC2.1^MTF2^ in the absence or presence of EPOP. Cells were harvested after 48 hours of transfection. Nuclear extracts were captured by the IgG resin, washed with the wash buffer (50 mM Tris-HCl, pH 8.0, 150 mM NaCl, 0.1% NP40, 2mM DTT, and 10% Glycerol), and released by the TEV cleavage or the TEV plus 3C combined cleavage overnight at 4°C. The eluted proteins were bound to the anti-FLAG resin and washed 3 times with the same wash buffer. The bound proteins were released by the wash buffer supplemented with 1 mg/ml FLAG peptide. The dimerization efficiency was assessed based on the anti-FLAG and anti-Myc signals in Western blot.

### EMSA

For the DNA^Lhx6^ binding assay, the 100 bp CGI^Lhx6^ DNA was amplified from mouse genomic DNA through PCR with fluorescein-labeled primers. CGI^Lhx6^ DNA (0.2nM) was incubated with purified PRC2.1^MTF2^ or PRC2.1^MTF2–EPOP^ (2-fold serial dilution starting from 1 μM) in a 20 μl reaction system, also containing 20mM Tris-HCl, pH 7.5, 100 mM KCl, 10% Glycerol, 1 mM DTT, 0.05% NP40, and 0.2 mg/ml yeast tRNA, for 30 min on ice. Samples were next loaded onto 4% native polyacrylamide gel (acrylacrylamide/bis 60:1) and resolved by electrophoresis in 1× TGE buffer (25 mM Tris, 190 mM Glycine, and 1 mM EDTA) at 100 V for 1 h on ice. The native gel was imaged by a Typhoon scanner. For the nucleosome binding assay, mononucleosome was assembled with the “601” DNA sequence using the standard salt dialysis method. Nucleosome (0.5 nM) was incubated with PRC2.1^MTF2^, PRC2.1^MTF2–EPOP^, or PRC2.1^MTF2–EPOP–Elongin^ ^BC^ (2-fold serial dilution starting from 1 μM) in a 20 μl reaction system, also containing 10 mM Tris-HCl, pH 8.0, 100 mM NaCl, and 10% Glycerol, on ice for 30 minutes. The mixture was resolved on a 4% native polyacrylamide gel in 1× TGE buffer at 100 V for 1 h on ice. The native gel was stained with SYBR Gold. The binding assays were repeated 3 times for each complex. Gel band quantification was performed using Image J, and the dissociation constant (Kd) was calculated by fitting binding curves in GraphPad Prism.

### Biotinylated DNA pull-down assay

The biotinylated CGI^Lhx6^ DNA was prepared as previously reported with minor modifications ^32^. Briefly, a PCR primer was labeled with biotin and contained an ECoRV cleave site before the CGI^Lhx6^ sequence. 1 μg PCR-amplified biotinylated CGI^Lhx6^ was incubated with avidin beads for 2 hours at 4°C and subjected to 3× wash with the wash buffer (50 mM Tris-HCl, pH 8.0, 150 mM NaCl, 0.1% NP40, 2mM DTT, and 10% Glycerol) to remove the unbound DNA. Nuclear extracts from WT or EPOP KO E14 mESCs were mixed with the DNA-bound beads and incubated at 4°C for 4 hours. Next, the beads were washed 3 times with the wash buffer and 2 times with the NEB CutSmart buffer. 1 μl ECoRV was added to the resin and incubated at 16°C overnight to cleave the DNA and release the bound protein.

### Expression and purification of the SUZ12(N)-RBBP4 binary complex

Human SUZ12 (residues 76-545τι146-155) and human RBBP4 (residues 1-425) were transiently co-expressed in HEK293T cells with the following plasmids: pHEK293Ultra- kozak-ProteinA-TEV-SUZ12 (residues 76-545 Δ146-155) and pHEK293Ultra-His6- thrombin-RBBP4 (residues 1-425). Transfection was performed when HEK293T cells reached 70% confluence, and transfected cells were harvested after 60 hours. For purification, the cell pellet was resuspended in lysis buffer (50 mM Tris-HCl, pH 8.0, 150 mM NaCl, 2 mM DTT, and 5% Glycerol) and sonicated to lyse the cells. The cell lysate was clarified by centrifugation at 24,600 x g for 30 minutes. An affinity pull-down was performed by incubating the cell lysate with IgG resin on a nutator at 4°C for 2 hours.

The IgG resin was washed with the wash buffer 1 (20 mM Tris-HCl, pH 8.0, 500 mM NaCl, 2 mM DTT, 5% Glycerol, and 0.1% NP-40), wash buffer 2 (20 mM Tris-HCl, pH 8.0, 1 M NaCl, 2 mM DTT, 5% Glycerol), and wash buffer 3 (20 mM Tris-HCl, pH 8.0, 150 mM NaCl, 2 mM DTT, 5% Glycerol). TEV protease was added for overnight cleavage. The binary complex released from the IgG resin was eluted, concentrated, and further purified by size-exclusion chromatography (SEC) using a Superdex200 10/300 GL column in the SEC buffer (20 mM Tris-HCl, pH 8.0, 150 mM NaCl, 2 mM DTT). Protein from peak fractions was collected, concentrated, aliquoted, flash-frozen, and stored in a -80°C freezer.

### Expression and Purification of the EPOP fragment

EPOP was expressed in the Rosetta2 *E. coli* strain. Plasmid pET28a-His6-SUMO-TEV- EPOP (residues 311-379)-FLAG was transformed into Rosetta2 competent cells, plated on LB agar plates containing kanamycin and chloramphenicol, and incubated at 37°C overnight. The following day, the colonies on plates were resuspended by 20 ml of LB medium, and the entire 20 ml cell suspension was inoculated into 1 liter of LB medium containing kanamycin and chloramphenicol. The cell culture was cultured at 37°C in a 250 rpm shaker until O.D.600 reached 0.6. Protein expression was induced using 0.5 mM IPTG at 20°C for 16 hours. For purification, the cell pellet was resuspended in the lysis buffer (100 mM HEPES, pH 7.4, 300 mM NaCl, 2 mM BME, 20 mM Imidazole, and 5% Glycerol) and sonicated to lyse the cells. The cell lysate was clarified by centrifugation at 24,600 x g for 30 minutes. The clarified cell lysate was mixed with Ni-NTA resin and incubated on a nutator at 4°C for 1 hour. The Ni-NTA resin was washed with wash buffer 1 (50 mM HEPES, pH 7.4, 1 M NaCl, 2 mM BME, 20 mM Imidazole, 0.1% NP-40, and 5% Glycerol) and wash buffer 2 (50 mM HEPES, pH 7.4, 1 M NaCl, 2 mM BME, 20 mM Imidazole, and 5% Glycerol). The resin was collected in a filter column, and 10 bed volumes of elution buffer (20 mM HEPES, pH 7.4, 150 mM NaCl, 2 mM BME, 250 mM Imidazole, and 10% Glycerol) were applied to the Ni-NTA resin.

SUMO protease was added to the eluted protein for overnight digestion. The next day, the digested protein solution was incubated with magnetic anti-FLAG resin on a nutator at 4°C for 2 hours. The anti-FLAG resin was washed using wash buffer 1 (20 mM Tris-HCl, pH 8.0, 500 mM NaCl, 2 mM BME, 5% Glycerol, and 0.1% NP-40) and wash buffer 2 (20 mM Tris, pH 8.0, 1 M NaCl, 2 mM BME, and 5% Glycerol). EPOP was eluted using the elution buffer containing the FLAG peptide (20 mM Tris-HCl, pH 8.0, 1 M NaCl, 2 mM BME, 5% Glycerol, and 0.5 mg/ml FLAG peptide). The protein was concentrated, aliquoted, flash-frozen, and stored at a -80°C freezer.

### Expression and Purification of the PHF19 fragment

PHF19 (residues 531-580) was expressed in Rosetta2 strain. Plasmid pGex4T1-GST-TEV-PHF19 (residues 531-580) was transformed into Rosetta2 competent cells, plated on an LB agar plate with ampicillin and chloramphenicol, and incubated at 37°C overnight. The next morning, the plate was washed with 20 ml of LB medium, and the entire 20 ml cell suspension was inoculated into 6 liters of LB medium with ampicillin and chloramphenicol. The cell culture was grown at 250 rpm in a 37 °C shaker until the O.D.600 reached 0.8. Protein expression was induced using 0.5 mM IPTG at 20°C for 16 hours. For purification, the cell pellet was resuspended in the lysis buffer (20 mM Tris-HCl, pH 8.0, 1M NaCl, 5 mM DTT, and 5% Glycerol) and sonicated to lyse the cells. The cell lysate was clarified by centrifugation at 24,600 x g for 30 minutes. The cell lysate was mixed with GST resin and incubated on a nutator at 4°C for 1 hour. The GST resin was washed with wash buffer 1 (20 mM Tris-HCl, pH 8.0, 1 M NaCl, 5 mM DTT, 0.1% NP-40, and 5% Glycerol) and wash buffer 2 (20 mM HEPES, pH 7.4, 1 M NaCl, 5 mM DTT, and 5% Glycerol). The resin was collected in a filter column, and 10 bed volumes of elution buffer (20 mM HEPES, pH 7.4, 1 M NaCl, 5 mM DTT, 25 mM Glutathione, and 5% Glycerol) were applied to the GST resin for elution. TEV protease was added to the protein elution to cleave the GST tag overnight. The resulting protein mixture was mixed with guanidine HCl powder to a final concentration of 7M and incubated at 4°C for 1 hour. The denatured protein mixture was dialyzed against a urea buffer (20 mM sodium acetate, pH 5.2, 10 mM NaCl, 5 mM BME, and 7M urea) using a 3.5 kD cutoff dialysis bag. After 4 hours, the dialysis bag was transferred to a new urea buffer, and the dialysis continued overnight. After dialysis, the protein mixture was loaded onto an SP column with low salt buffer (10 mM sodium acetate, pH 5.2, 10 mM NaCl, and 5 mM BME). It was washed with low salt buffer and then directly eluted using high salt buffer (10 mM Sodium acetate, pH 5.2, 1 M NaCl, and 5 mM BME) to obtain the highly concentrated PHF19 fragment. Finally, the PHF19 fragment was dialyzed against water containing 5 mM BME overnight to remove urea. The protein solution was aliquoted, flash-frozen, and kept at -80°C.

### Reconstitution of the heterotetrameric complex and crystal structure determination

The SUZ12(N)–RBBP4 binary complex was mixed with the PHF19 fragment at a molar ratio of 1:1.2 and incubated on ice for 1 hour. The EPOP fragment was next added such that a molar ratio of SUZ12(N)–RBBP4:PHF19:EPOP became 1:1.2:2.0, and the protein complex was incubated on ice for another 1 hour. The protein complex was purified using a Superdex200 10/300 GL column in the SEC buffer (20 mM Tris-HCl, pH 8.0, 100 mM NaCl, and 2 mM DTT). Peak fractions were collected, concentrated, aliquoted, and flash-frozen at -80°C freezer. Crystals of the heterotetrameric complex were obtained using hanging drop vapor diffusion against a reservoir buffer (100 mM sodium citrate, pH 5.5, 200 mM sodium acetate, 9% PEG4000) at 20°C for 1-2 weeks. The crystals were harvested and frozen in a cryo-protecting buffer (100 mM sodium citrate, pH 5.5, 200 mM sodium acetate, 12% PEG4000, 20% glycerol). Single crystal diffraction data were collected at a synchrotron light source and processed by HKL- 2000 ^60^. The structure was solved by molecular replacement in Phaser using PDB 5WAK as the search model ^61^. The structural model was built in Coot and refined in autoBUSTER ^62,63^. The structure refinement statistics were generated in Phenix ^64^. Structural images were rendered in PyMOL ^65^.

---

Table S1 Diffraction data collection and structure refinement statistics

Table S2 RNA-seq and differential gene expression

Table S3 Differential chromatin binding in EpiLCs

Table S4 PRC2.1 repressed EPOP-maintained genes in EpiLCs

