## Supplemental Figures for "EPOP Restricts PRC2.1 Targeting to Chromatin by Directly Modulating Enzyme Complex Dimerization"

**Fig. S1** **Gong L. et al.**

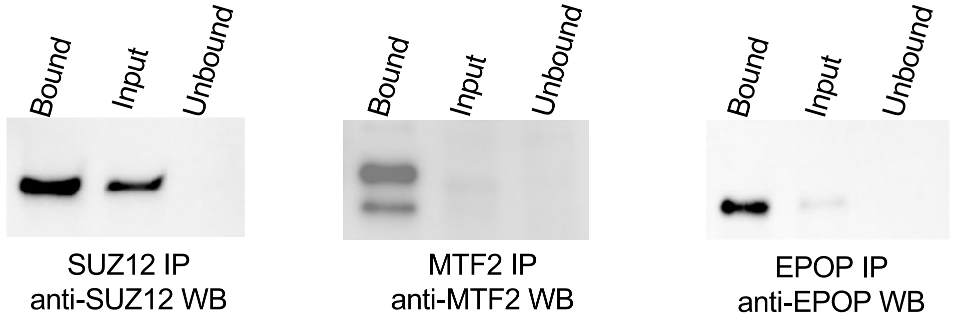

**Fig. S1 Immunodepletion of endogenous PRC2 components in mESCs.**

Home-made antibody beads were tested for the efficiency of the immunodepletion of SUZ12, MTF2, and EPOP.

### Fig. S2

Gong L. et al.

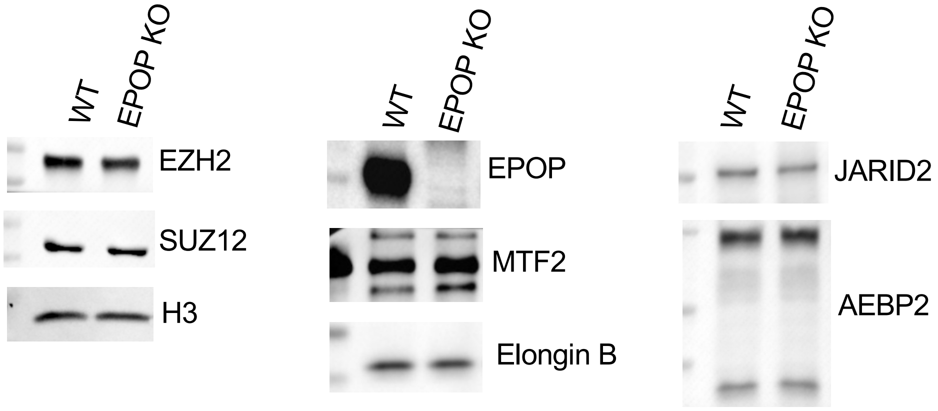

**Fig. S2 Expression level of PRC2 components in WT and EPOP KO mESCs.**

Western blot was used to compare the expression levels of selected PRC2 components, EZH2, SUZ12, EPOP, MTF2, Elongin B, JARID2, and AEBP2.

**Fig. S3** **Gong L. et al.**

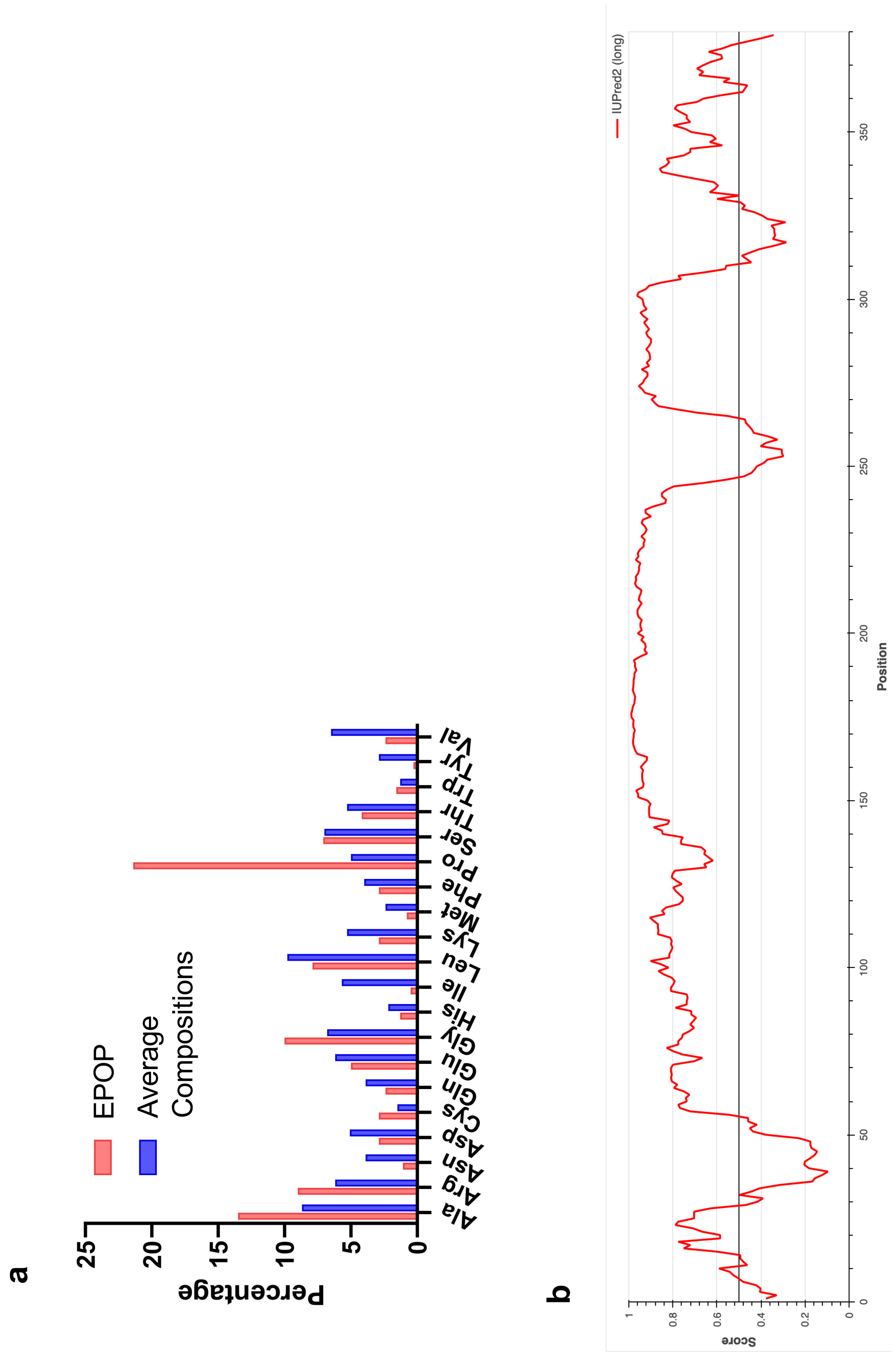

**Fig. S3 Intrinsically disordered EPOP protein sequence.**

**a**, Amino acid composition of EPOP compared to average proteins. Information on average proteins was based on Proteopedia (https://proteopedia.org/wiki/index.php/Amino_acid_composition).

**b**, Intrinsically disordered regions of EPOP. The prediction was made using the IUPred2 server (https://iupred2a.elte.hu).

**a**

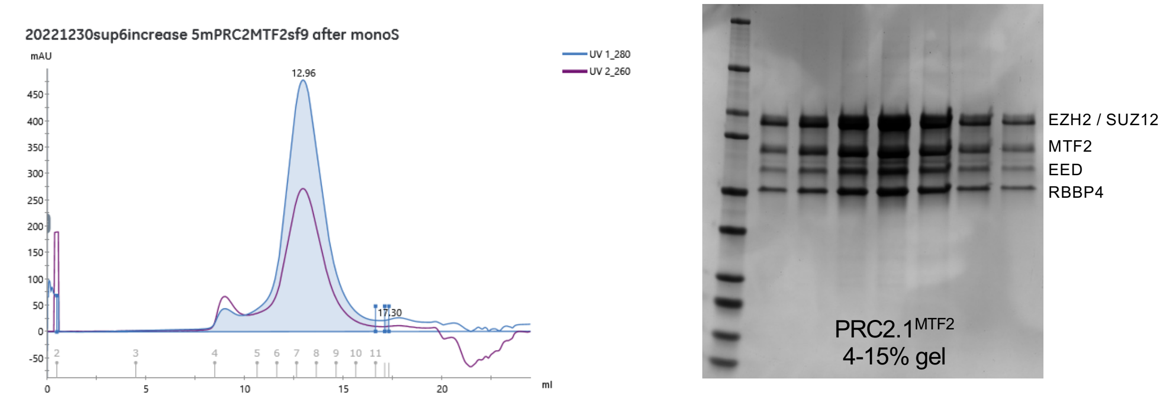
20221230sup6increase 5mPRC2MTF2sf9 after monoS
mAU
UV 1_280
UV 2_260
12.96
17.30
ml
EZH2 / SUZ12
MTF2
EED
RBBP4
PRC2.1MTF2
4-15% gel

**b**

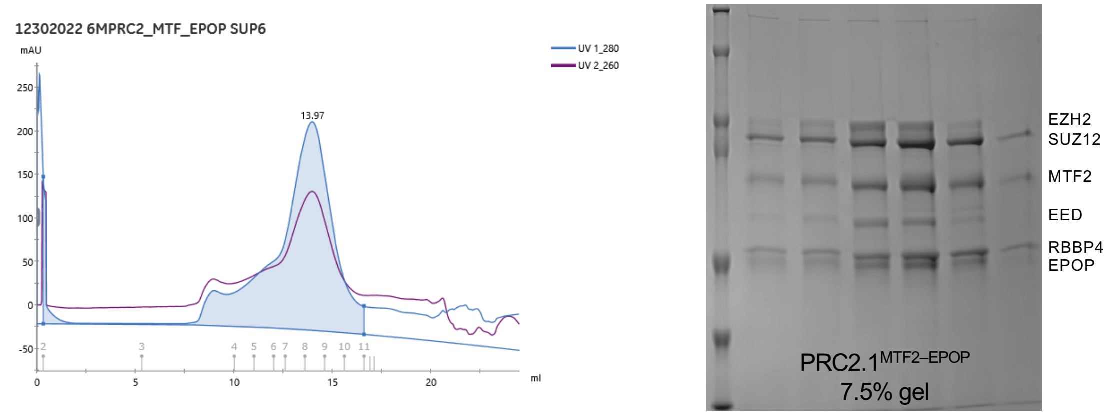
12302022 6MPRC2_MTF_EPOP SUP6
mAU
UV 1_280
UV 2_260
13.97
ml
EZH2
SUZ12
MTF2
EED
RBBP4
EPOP
PRC2.1MTF2–EPOP
7.5% gel

**c**

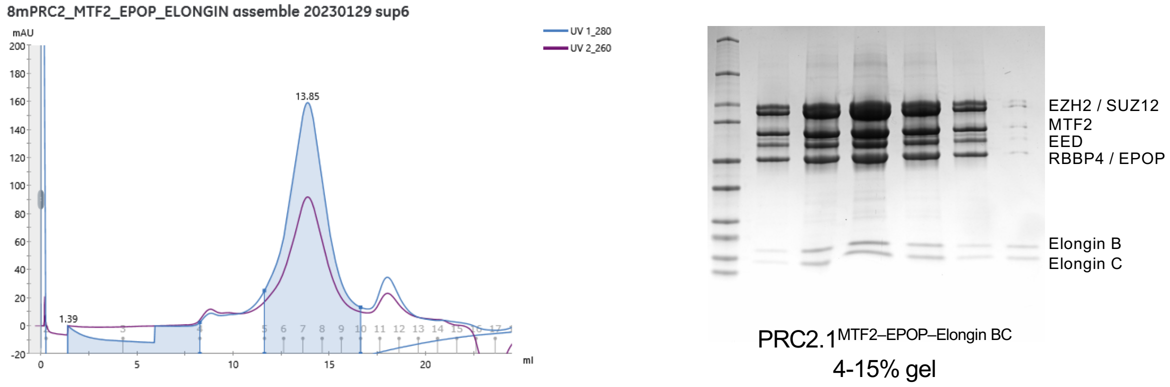
8mPRC2_MTF2_EPOP_ELONGIN assemble 20230129 sup6
mAU
UV 1_280
UV 2_260
13.85
1.39
ml
EZH2 / SUZ12
MTF2
EED
RBBP4 / EPOP
Elongin B
Elongin C
PRC2.1MTF2–EPOP–Elongin BC
4-15% gel

**Fig. S4 Purification of PRC2.1 holocomplexes.**

SEC profiles and corresponding SDS-PAGE gels are shown for **a.** PRC2.1$^{MTF2}$, **b.** PRC2.1$^{MTF2–EPOP}$, and **c.** PRC2.1$^{MTF2–EPOP–Elongin\ BC}$.

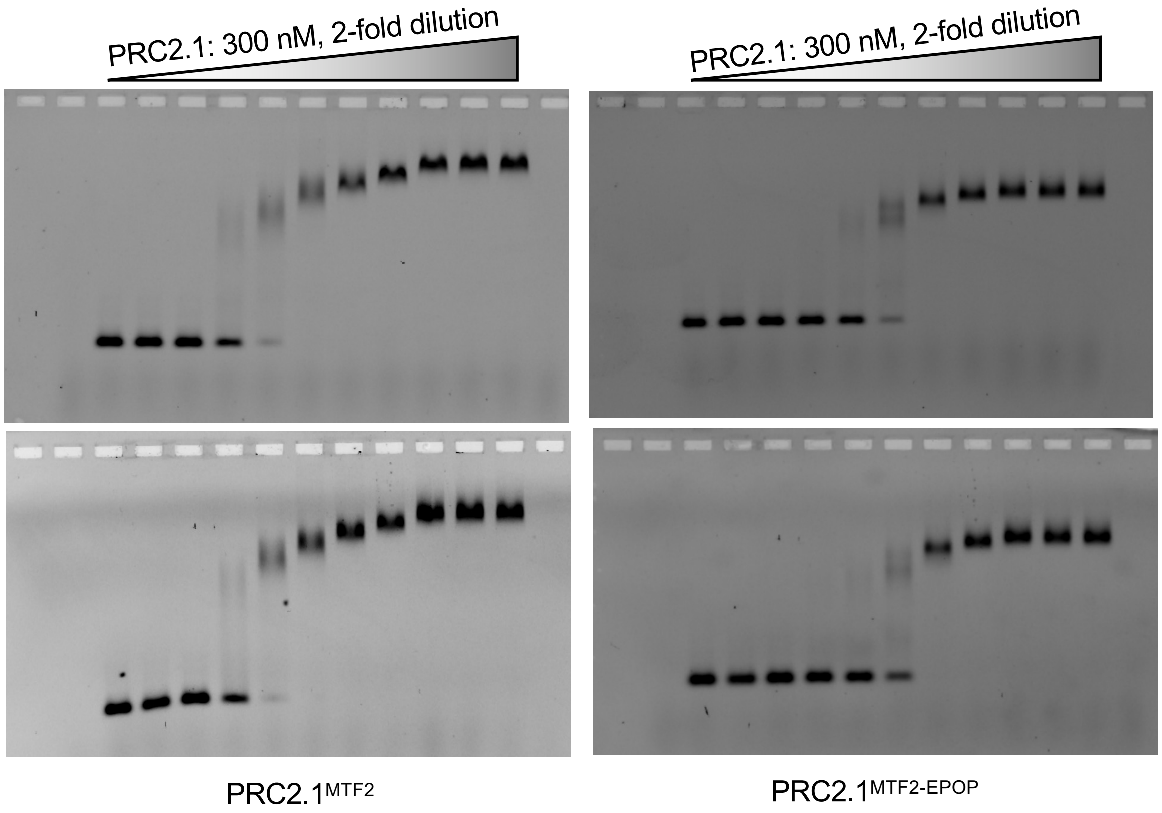
PRC2.1: 300 nM, 2-fold dilution
PRC2.1: 300 nM, 2-fold dilution
PRC2.1$^{MTF2}$
PRC2.1$^{MTF2\text{-}EPOP}$

**Fig. S5 DNA binding EMSA.**

Replicates 2 and 3 of $CGI^{\textit{Lhx6}}$ DNA binding by $PRC2.1^{MTF2}$ and $PRC2.1^{MTF2–EPOP}$ are shown.

**Fig. S6** **Gong L. et al.**

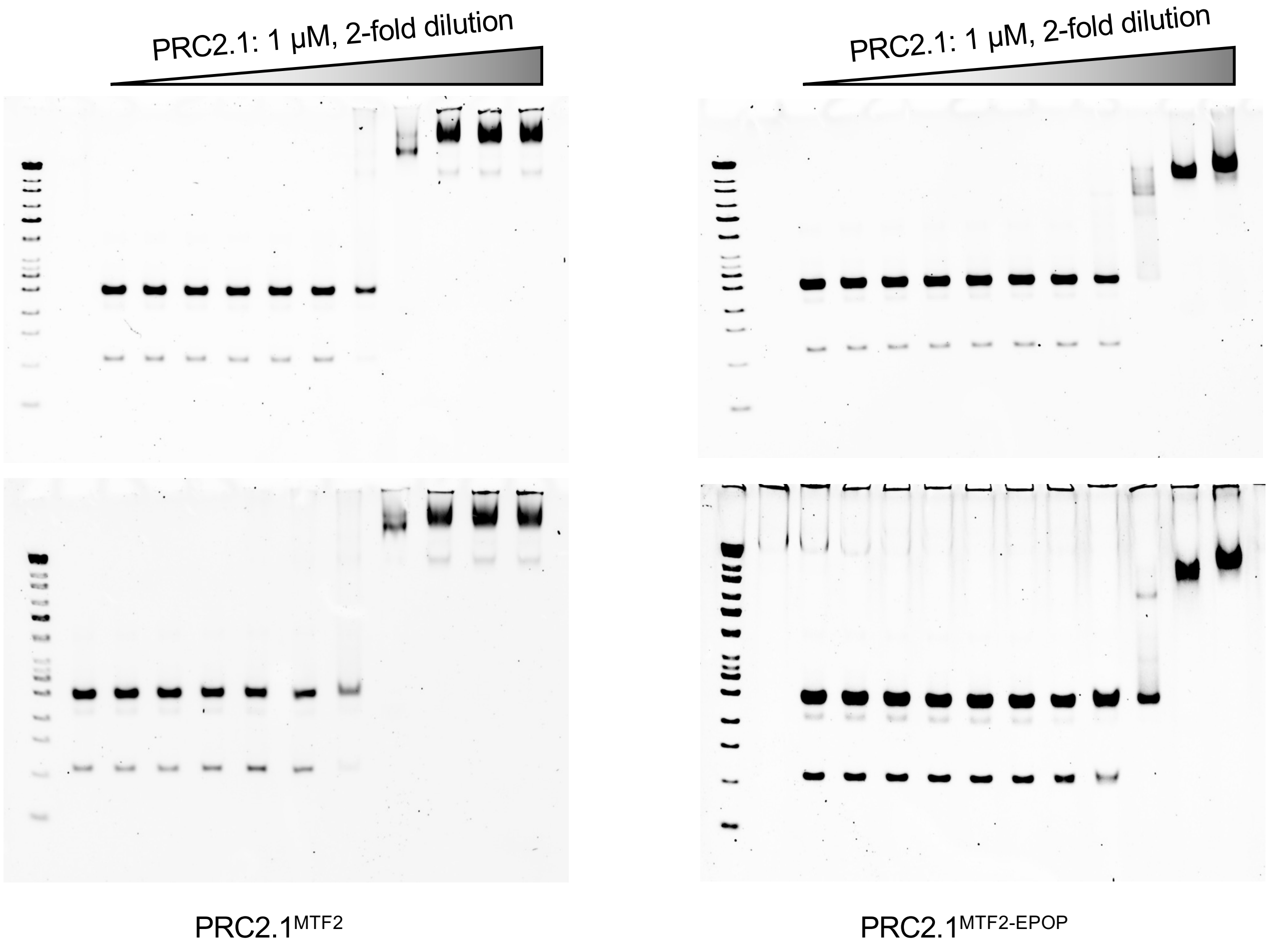

**Fig. S6 Nucleosome binding EMSA for PRC2.1$^{MTF2}$ and PRC2.1$^{MTF2–EPOP}$.**

Replicates 2 and 3 of nucleosome binding by PRC2.1$^{MTF2}$ and PRC2.1$^{MTF2–EPOP}$ are shown.

Fig. S7

Gong L. et al.

a

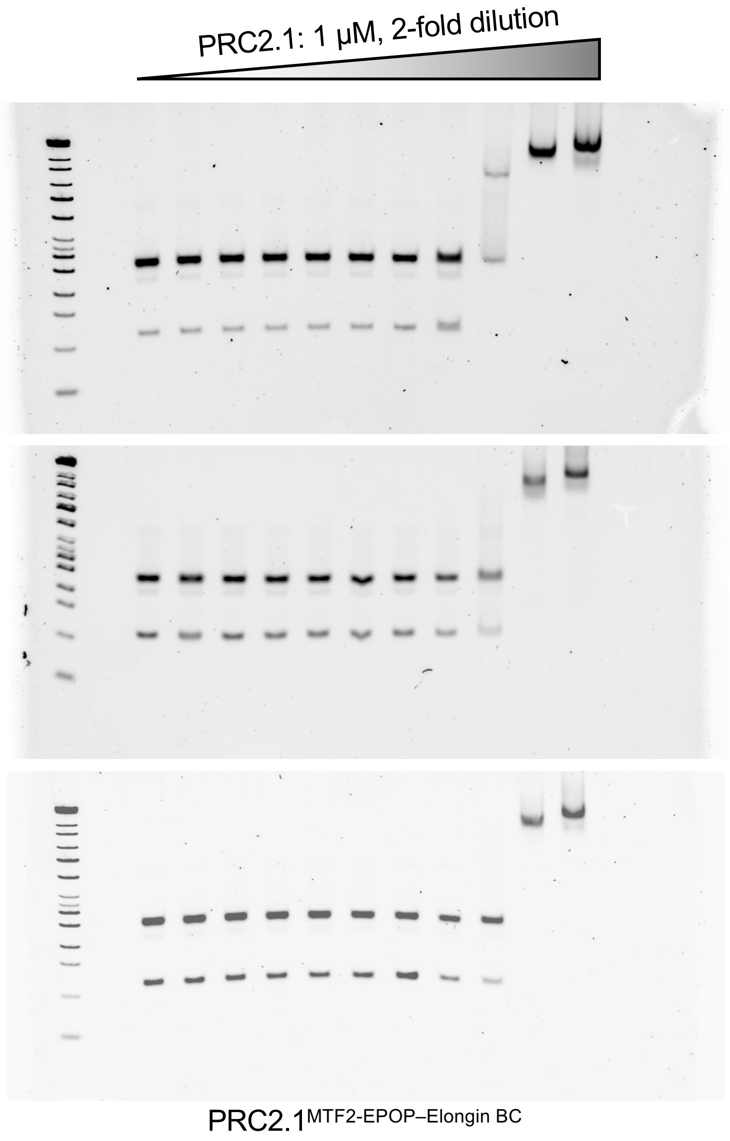


b

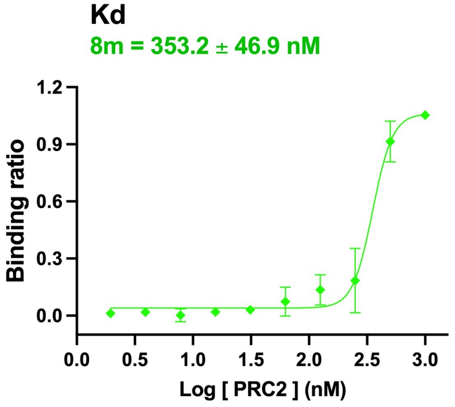

**Fig. S7 Nucleosome binding EMSA for PRC2.1$^{\text{MTF2–EPOP–Elongin BC}}$.**

**a.** Three replicates of nucleosome binding by PRC2.1$^{\text{MTF2–EPOP–Elongin BC}}$ are shown.

**b.** The binding affinity was derived from the binding curve.

### Fig. S8

Gong L. et al.

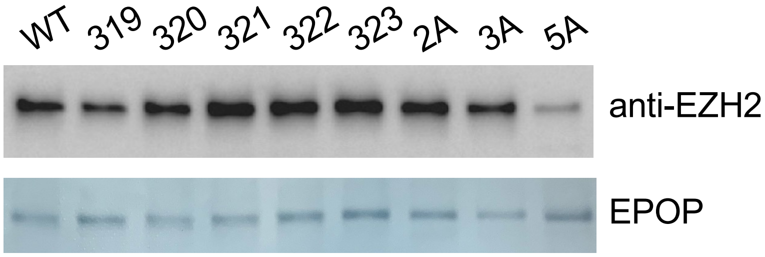

**Fig. S8 Co-IP assay of EPOP.**

$His_6$-SUMO-Protein A-TEV-EPOP-Myc WT and mutants were transiently expressed in HEK293T cells. IgG beads were used to pull down EPOP proteins, and bound endogenous EZH2 was released by TEV cleavage and detected by Western blot. The bait EPOP protein was detected on the PVDF membrane using amido black. 319, F319A; 320, P320A; 321, C321A; 322, P322A; 323, P323A; 2A, P322A/P323A; 3A, P320A/C321A/P322A; 5A, F319A/P320A/C321A/P322A/P323A.

### Fig. S9

Gong L. et al.

**a**

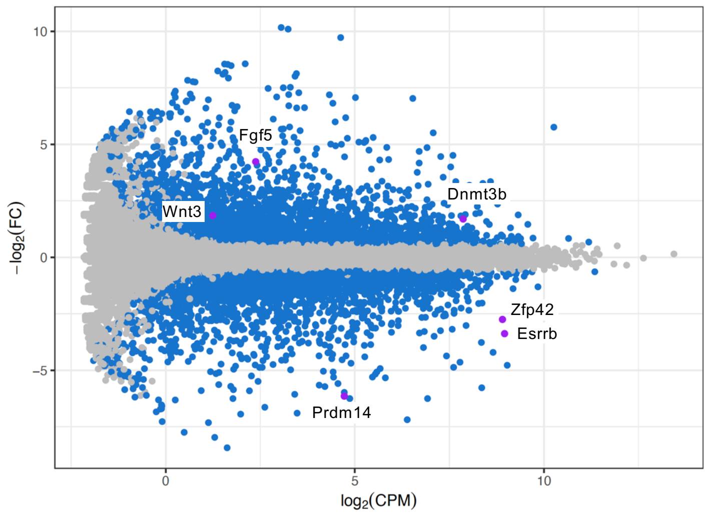


ESC-to-EpiLC, WT

**b**

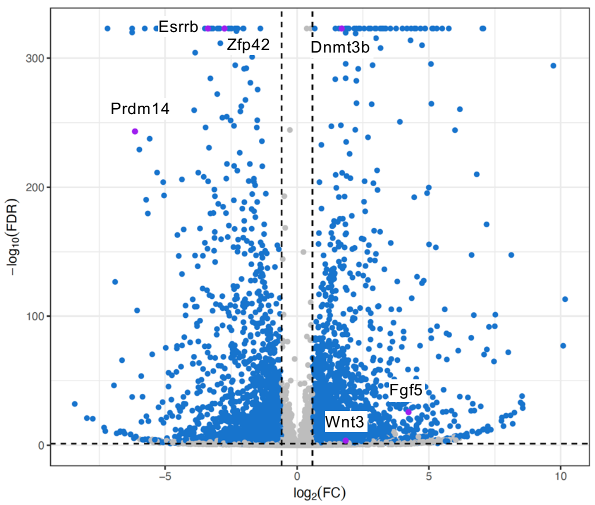


ESC-to-EpiLC, WT

**c**

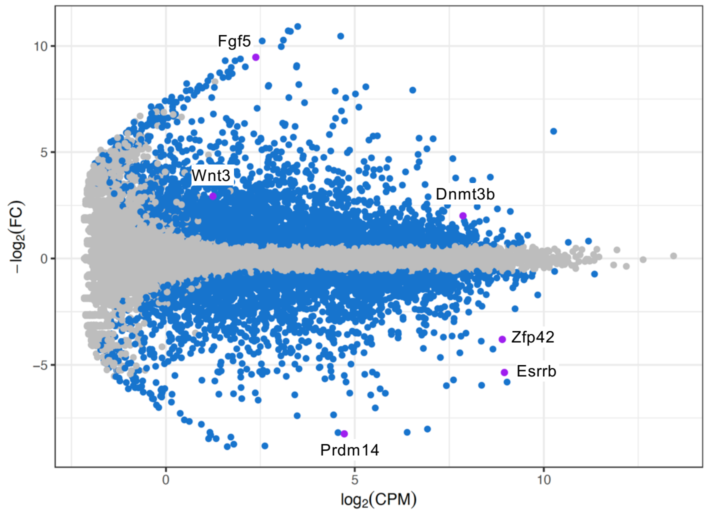


ESC-to-EpiLC, D5

**d**

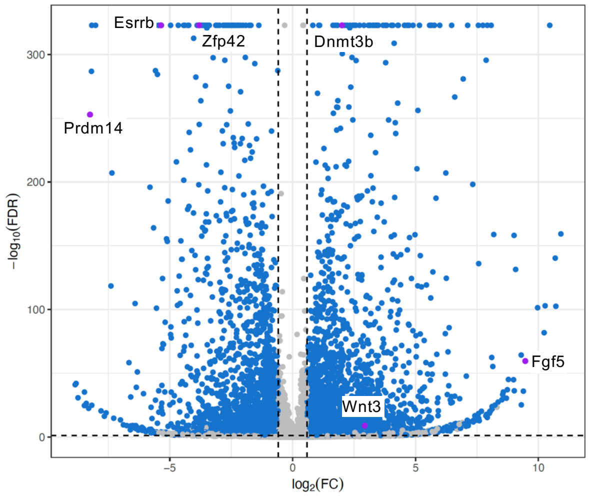


ESC-to-EpiLC, D5

**Fig. S9 MA and volcano plots of the EpiLC differentiation.**

**a**, MA plot of the differential gene expression during the EpiLC differentiation of the EPOP$^{WT}$ mESCs.

**b**, Volcano plot of **a**.

**c**, MA plot of the differential gene expression during the EpiLC differentiation of the EPOP$^{D5}$ mESCs.

**d**, Volcano plot of **c**.

Naïve and EpiLC mark genes are highlighted by purple dots and labeled.

### Fig. S10

Gong L. et al.

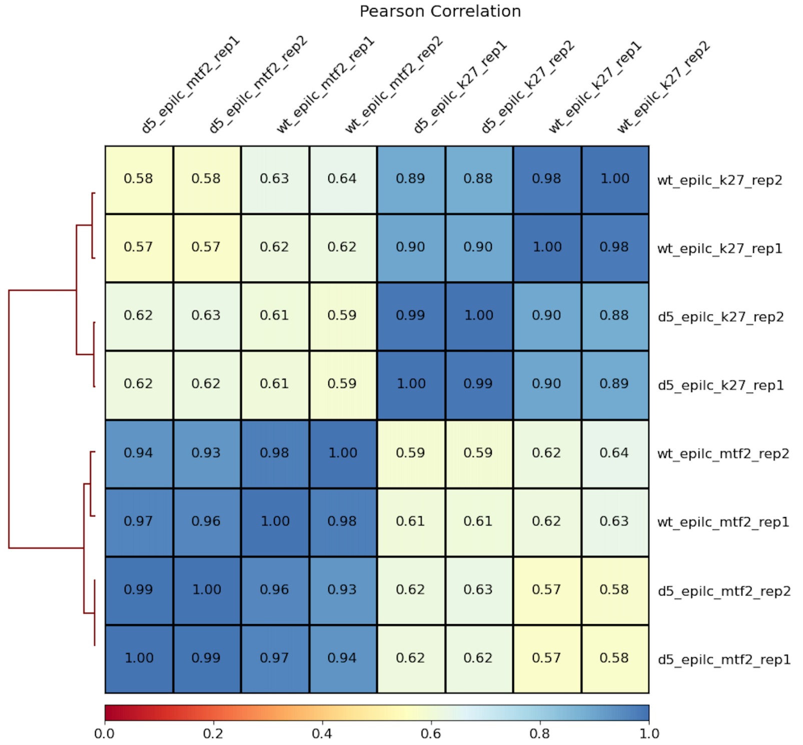

**Fig. S10 Correlation of ChIP-seq data sets.**

The Consistency of ChIP-seq replicates was checked by Pearson correlation.

### Fig. S11

Gong L. et al.

a

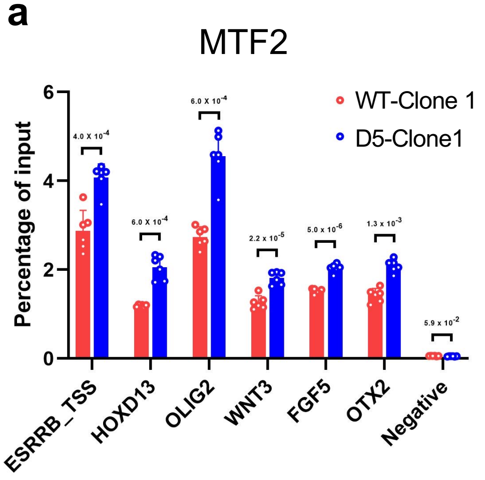


b

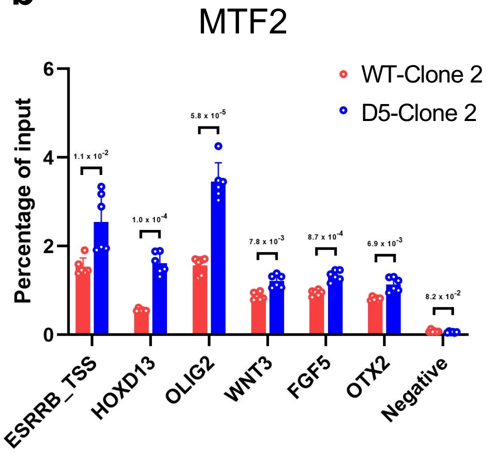

**Fig. S11 ChIP-qPCR of the MTF2 enrichment**

Two independent pairs of WT and D5 clones were differentiated and checked by ChIP-qPCR. The Clone 1 pair were used for ChIP-seq.
**a**, MTF2 enrichments at selected PRC2 targets in the Clone 1 pair were compared.

**b**, MTF2 enrichments at selected PRC2 targets in the Clone 2 pair were compared.

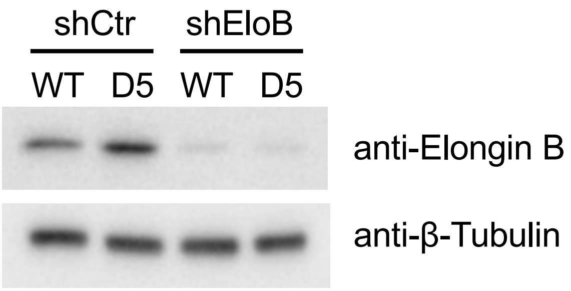
shCtr
shEloB
WT
D5
WT
D5
anti-Elongin B
anti-β-Tubulin

**Fig. S12 Elongin B KD.**

The efficiency of the control and Elongin B stable KD with shRNAs was checked by Western blot.

### Fig. S13

Gong L. et al.

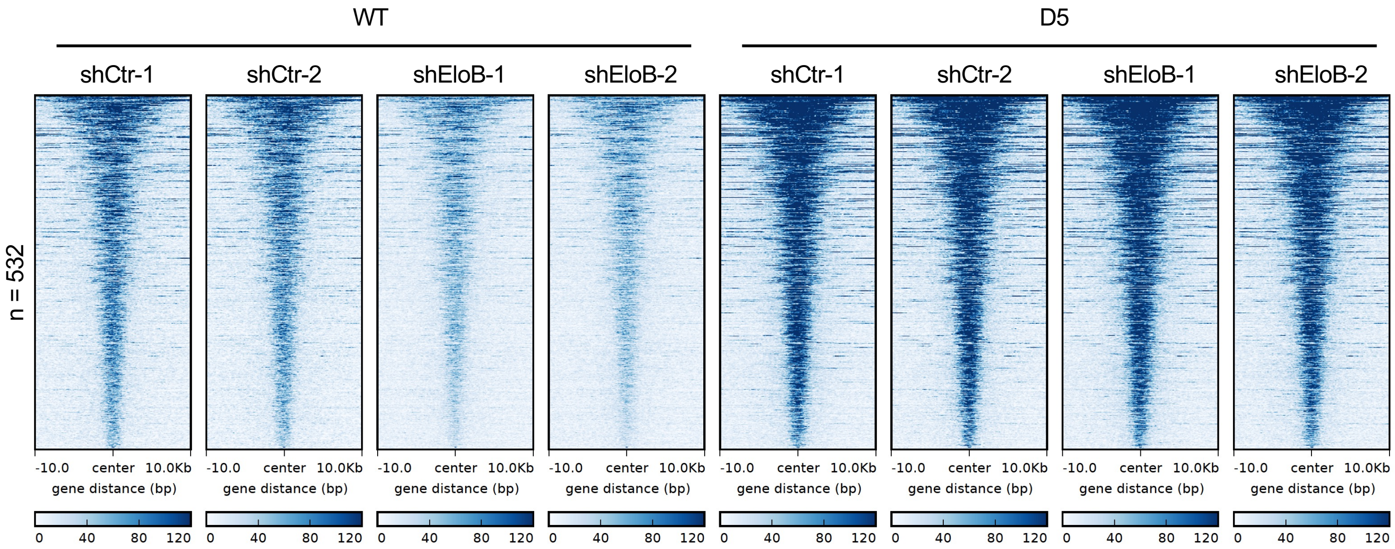

**Fig. S13 Positive role of Elongin BC on MTF2 targeting in EpiLCs.**

A set of 532 consensus MTF2 binding sites lost signals upon Elongin B KD in the $EPOP^{WT}$ EpiLCs but not in the $EPOP^{D5}$ EpiLCs.

**Fig. S14** **Gong L. et al.**

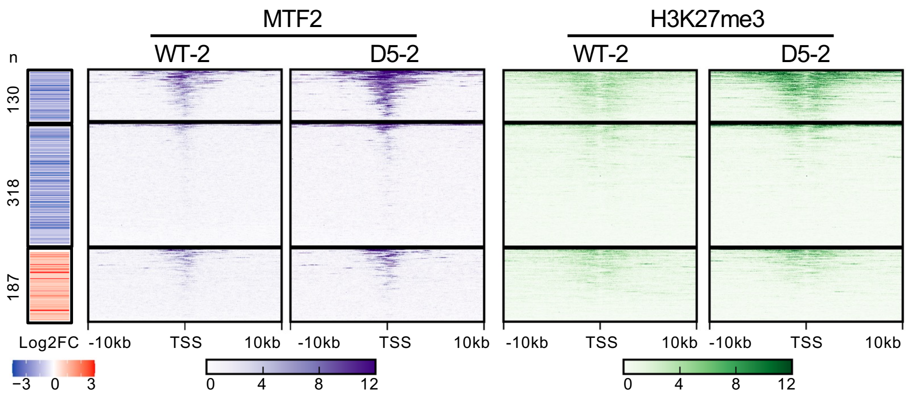

**Fig. S14 Correlation of RNA-seq and ChIP-seq.**

ChIP-seq replicate data shows a correlation between gene expression and MTF2 chromatin binding around the TSS for a fraction of downregulated genes. See the main figure for details.

**Fig. S15**

**Gong L. et al.**

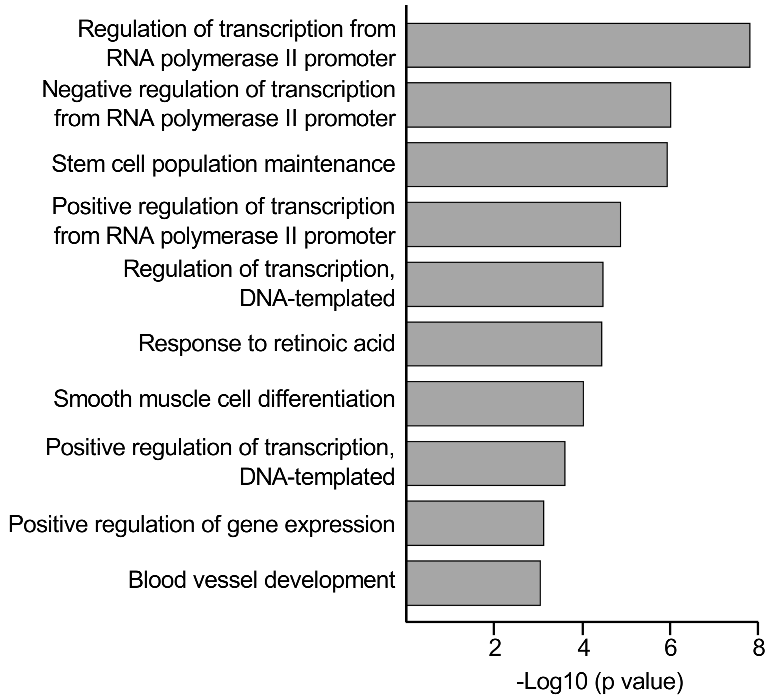

**Fig. S15 Gene ontology analysis.**

The 130 PRC2.1 repressed, EPOP-maintained genes in EpiLCs were subjected to gene ontology analysis for associated biological processes.  Top 10 hits are summarized.

**Supplementary Table 1. Diffraction Data Collection and Structure Refinement Statistics**

| **Data collection** | |
|---|---|
| Wavelength (Å) | 0.979 |
| Resolution range (Å) | 48.4-2.7 (2.8-2.7) |
| Space group | R32 |
| a, b, c (Å) | 148.8,148.8, 290.7 |
| $\alpha$, $\beta$, $\gamma$ (°) | 90.0, 90.0, 120.0 |
| $R_{merge}$ (%) | 13.9 (154.8) |
| $R_{pim}$ (%) | 3.2 (44.0) |
| Mean I/$\sigma$ | 25.7 (1.6) |
| $CC_{1/2}$(%) | 99.7 (59.8) |
| Completeness (%) | 99.8 (98.3) |
| Redundancy | 19.1 (13.3) |
| **Refinement** | |
| Number of reflections | 34574 |
| $R_{work}$/$R_{free}$ | 0.209 /0.253 |
| Number of non-hydrogen atoms | 6163 |
| RMS(bonds) | 0.012 |
| RMS(angles) | 1.62 |
| Ramachandran favored (%) | 96.21 |
| Ramachandran allowed (%) | 3.38 |
| Ramachandran outliers (%) | 0.41 |
